# A gas-and-brake mechanism of bHLH proteins modulates shade avoidance in *Arabidopsis thaliana*

**DOI:** 10.1101/2020.05.07.082735

**Authors:** Sara Buti, Chrysoula K. Pantazopoulou, Valérie Hoogers, Kasper van Gelderen, Emilie Reinen, Ronald Pierik

## Abstract

Plants detect proximity of competitors through reduction in the ratio between red and far red light triggering the shade avoidance syndrome, which includes accelerated shoot elongation and early flowering. Shade avoidance is regulated through PHYTOCHROME INTERACTING FACTORs (PIFs), a group of bHLH transcription factors. Another (b)HLH protein, KIDARI (KDR), which is non-DNA-binding, was identified in de-etiolation studies and proposed to interact with LONG HYPOCOTYL IN FAR-RED 1 (HFR1), a (b)HLH protein that inhibits shade avoidance. Here we establish novel roles of KDR in regulating shade avoidance and investigate how KDR regulates the shade avoidance network. We show that KDR is a positive regulator of shade avoidance and interacts with several negative growth regulators. We identify novel interactors using a combination of yeast two-hybrid screening and dedicated confirmations with bimolecular fluorescence complementation. We demonstrate that KDR is translocated primarily to the nucleus when coexpressed with these newly discovered interactors. A genetic approach confirmed that several of these novel interactions are indeed functional to shade avoidance in *Arabidopsis thaliana*, whereas we propose that KDR does not interact with HFR1 to regulate shade avoidance. Based on this, we propose that shade avoidance is regulated by a three-layered gas-and-brake mechanism of bHLH protein interactions, adding an additional layer of complexity to what was previously known.

**One-sentence summary:** KIDARI is a positive regulator of shade avoidance and part of a three-layer network of bHLH transcription factor interactions.

## INTRODUCTION

Plants harvest light energy during photosynthesis, especially, blue (B) (∼400-500 nm waveband) and red (R) (∼600-700 nm waveband) light, whilst mostly reflecting farred light (FR) (∼700-800 nm waveband). As a consequence, the ratio of R to FR is reduced by light reflected or transmitted through plant leaves and neighbors use this to detect the presence of nearby plants. Shade intolerant plants, such as *Arabidopsis thaliana*, respond to this lowered R:FR ratio with the shade avoidance syndrome (SAS). The main shade avoidance characteristics in *A. thaliana* are: hypocotyl, internode and petiole elongation, early flowering, and upward leaf movement called hyponasty (de Wit et al., 2016; Ballaré et al., 1991; de Wit et al., 2015; Galvão et al., 2019; Pantazopoulou et al., 2017). SAS is typical of most plants, including crops, and although it improves individual plant fitness, it may compromise total crop yield (Robson et al., 1996; Boccalandro et al., 2003). Contrary, shade tolerant species, such as those from forest understories, have developed alternative strategies to cope with shade conditions without investing in shade avoidance growth (Gommers et al., 2013, 2017; Molina-Contreras et al., 2019).

In an attempt to unravel the strategy of some species suppressing SAS, Gommers et al. (2017) previously described the contrasting shade-tolerant and intolerant responses of two selected *Geranium* species when exposed to low R:FR. In a transcriptome approach between these species, putative regulators of these two different responses were identified. One of these regulators is a basic helix-loop-helix (bHLH) protein-encoding gene, called *KIDARI (KDR)/PACLOBUTRAZOL RESISTANCE 6* (*PRE6*) (Gommers et al., 2017). The expression of *KDR* in *A. thaliana* has been shown to rely on functional PHYTOCHROME-INTERACTING FACTOR 4 (PIF4), PIF5 and PIF7 in both white light and low R:FR conditions (Gommers et al., 2017) but the precise functioning of KDR in shade avoidance responses is still to be elucidated.

The main mechanism by which KDR has been previously shown to regulate growth in dark versus monochromatic light is by acting as a cofactor. KDR does not directly regulate gene transcription because it lacks the capability to bind DNA, but it is able to interfere with the action of other proteins. KDR cannot bind DNA directly because it misses specific amino acids (glutamic acid and arginine) in the basic domain, which are essential for DNA binding (Hyun and Lee, 2006). Its function is mainly determined by the HLH domain and through this domain, KDR was proposed to interact with LONG HYPOCOTYL IN FAR-RED 1 (HFR1) (Hong et al., 2013; Hyun and Lee, 2006), another non-DNA-binding (b)HLH protein. HFR1 is an established regulator of shade avoidance and binds to PIF proteins (Hornitschek et al., 2009), preventing them from activating the transcription of genes associated with SAS. HFR1 and PIF4 are both members of the bHLH transcription factor (TF) family, but while PIF4 is promoting SAS through the transcriptional activation of specific genes, HFR1 plays a negative role in SAS by suppressing PIF4 action through direct binding (Sessa et al., 2005; Hornitschek et al., 2009). It is proposed that the regulation of both positive and negative regulators upon shade exposure helps plants tune the intensity of their shade avoidance responses (Gommers et al., 2017; Sessa et al., 2005; de Wit et al., 2016). *PIF4* and other members of the same sub-family, such as *PIF1, PIF3, PIF5* and *PIF7*, are not transcriptionally upregulated in shade conditions but their proteins are stabilized (Al-Sady et al., 2006; Shen et al., 2007; Leivar et al., 2008; Li et al., 2012; Lorrain and Fankhauser, 2012). In standard light, phytochromes are active and through interaction with PIFs, lead to PIF inactivation and often degradation, whereas phytochromes are inactivated in shade, relieving the repression of PIF activity (Chen and Chory, 2011; Leivar and Quail, 2011). PIF proteins act as positive regulators of SAS primarily by promoting auxin synthesis, transport and response (Hornitschek et al., 2012; Li et al., 2012). Simultaneously, PIFs also promote the expression of several negative regulators, such as *HFR1, PHYTOCHROME RAPIDLY REGULATED 1* (*PAR1*) and *PAR2* (Sessa et al., 2005; Roig-Villanova et al., 2006), all non-DNA binding (b)HLH proteins. Thus, there is a high redundancy as well specification within the bHLH family in the regulation of SAS, allowing a highly flexible response that can integrate different environmental parameters.

In the present study, we establish the role of KDR in shade avoidance. Using established and novel *A. thaliana kdr* mutant and overexpression lines, we demonstrate that KDR is a positive regulator of low R:FR-induced hypocotyl elongation. *KDR* overexpression, in addition to promoting hypocotyl elongation, also stimulates primary root elongation, bolting and flowering. Using yeast two-hybrid (Y2H) and bimolecular fluorescence complementation (BiFC) approaches, we identify several novel interactors of KDR and show that they colocalize with KDR in the nucleus. Experiments on mutants and transgenics to modulate the expression levels of these putative interactors, combined with published knowledge about these genes, suggest that all KDR-interacting proteins found here are themselves negative regulators of low R:FR-induced hypocotyl elongation. We propose that KDR interaction with these growth suppressors disables them from interacting with their downstream targets, alleviating the restraint on shade avoidance.

## RESULTS

### KDR promotes shade avoidance response

We confirmed the upregulation of *KDR* in seedlings of *A. thaliana* exposed to a low R:FR treatment (R:FR = 0.2) in comparison to control condition (R:FR = 2) (Figure 1A). To further investigate the role of KDR in SAS we studied the response of *kdr* lines to low R:FR conditions. We measured the elongation of hypocotyls in seedlings and of petioles in rosette plants upon exposure to low R:FR. Hypocotyl elongation of a *KDR* knockout line *kdr-1* was reduced in low R:FR conditions, whereas the activation-tagged line *kdr-D*, which results in four times the insertion of the *CaMV 35S* promoter in the promoter region of *KDR*, displayed an exaggerated elongation (Figure 1B). When the same lines were tested for petiole elongation in rosette plants, a similar suppression of the response was found for *kdr-1*, but less severely. By contrast, the overexpressing line *kdr-D* showed no statistically significant difference with Col-0 wild type for low R:FR-induced petiole elongation. The petiole elongation responses between wild type and mutants were not dependent on day length (Figure 1C).

**Figure 1:**
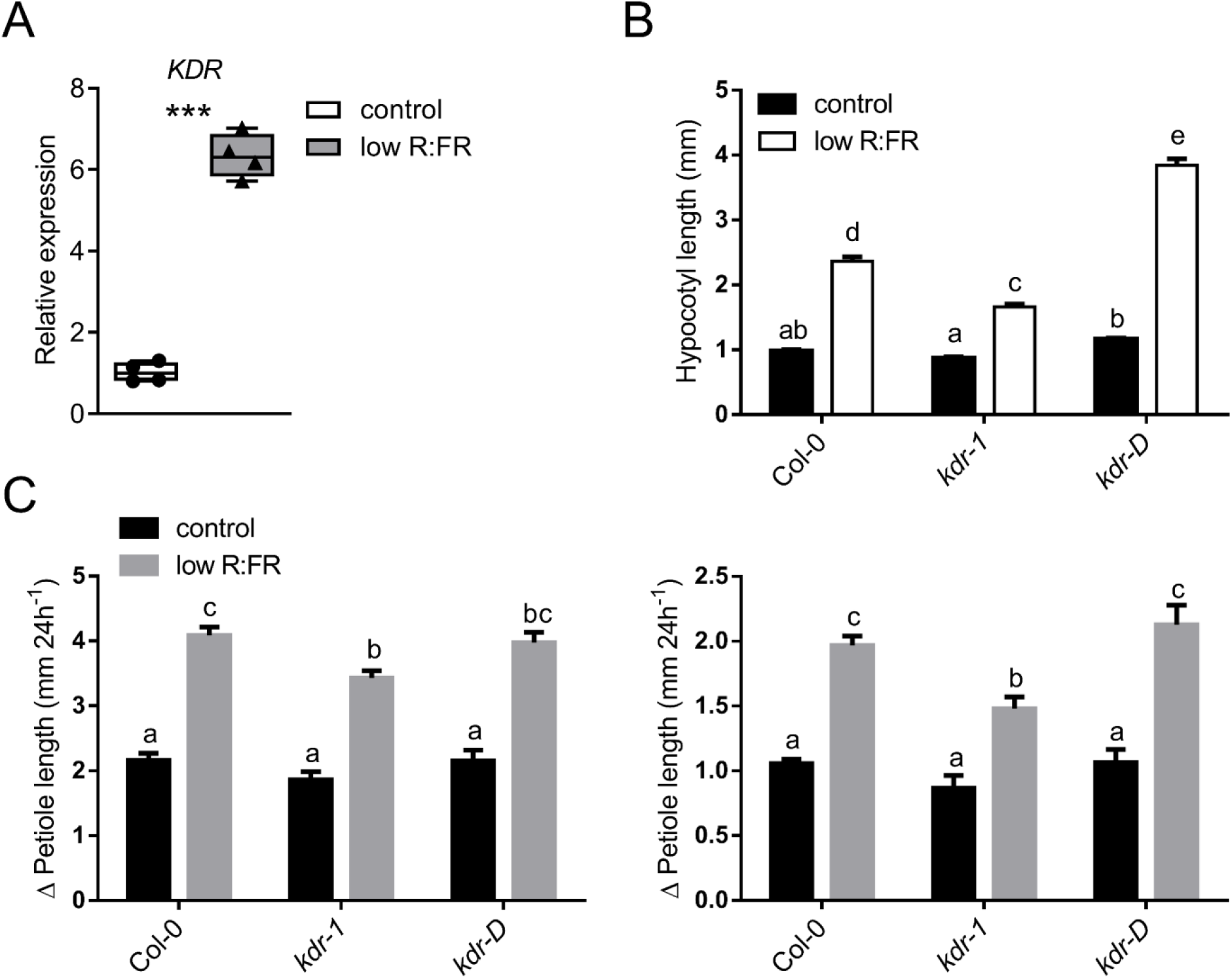
*A. thaliana kdr* mutants exhibit a deviating low R:FR response. (**A**) Relative expression of *KDR* determined by qRT-PCR in wild type Col-0 shoots grown in control white light condition (R:FR = 2) and low R:FR (R:FR = 0.2) for 90 minutes. Data represent mean ± SE, n = 4. Statistically significant difference is indicated by *** for p < 0.001, Student’s t-test. (**B**) Hypocotyl length (mm) of seedlings of *A. thaliana* wild type (Col-0), knockout line (*kdr-1*) or activation-tagged line (*kdr-D*) grown in control light condition (R:FR = 2) or low R:FR (R:FR = 0.2) after 5 days of light treatment. Data represent mean ± SE, n = 76. Different letters indicate statistically significant differences (2-way ANOVA with post-hoc Tukey test, p < 0.05). (**C**) Change in petiole length (Δ Petiole length) (mm) of *A. thaliana* wild type (Col-0) rosette plants, knockout line (*kdr-1*) or activation-tagged line (*kdr-D*) grown in control light condition (R:FR = 2) or low R:FR (R:FR = 0.2) after 24 h of light treatment. Plants are grown in short day (left) or in long day (right) conditions. Data represent mean ± SE, n = 10. Different letters indicate statistically significant differences (2-way ANOVA with post-hoc Tukey test, p < 0.05).

### Overexpression of *KDR* stimulates shade avoidance

We created novel lines overexpressing *KDR* in *A. thaliana* Col-0 background in order to have improved genetic material over the *kdr-D* activation tagging line that only mildly overexpresses *KDR.* We then used four homozygous independent lines to study their response to low R:FR treatment at the seedling stage. We found that most of the novel *35S:KDR* transgenic lines showed an even more exaggerated response than found for *kdr-D* and additionally displayed constitutive hypocotyl elongation in white light (Figure 2A). Interestingly, the variation in hypocotyl length between the independent transgenic lines correlated with variation in *KDR* overexpression levels (Figure 2B and 2C). We selected the two independent lines most strongly overexpressing *KDR* to continue. When looking more carefully at the phenotype of the selected lines, we observed that *KDR* overexpression increased elongation of most organs, including hypocotyl, petioles of cotyledons, petioles of true leaves and primary root (Figure 2D).

**Figure 2:**
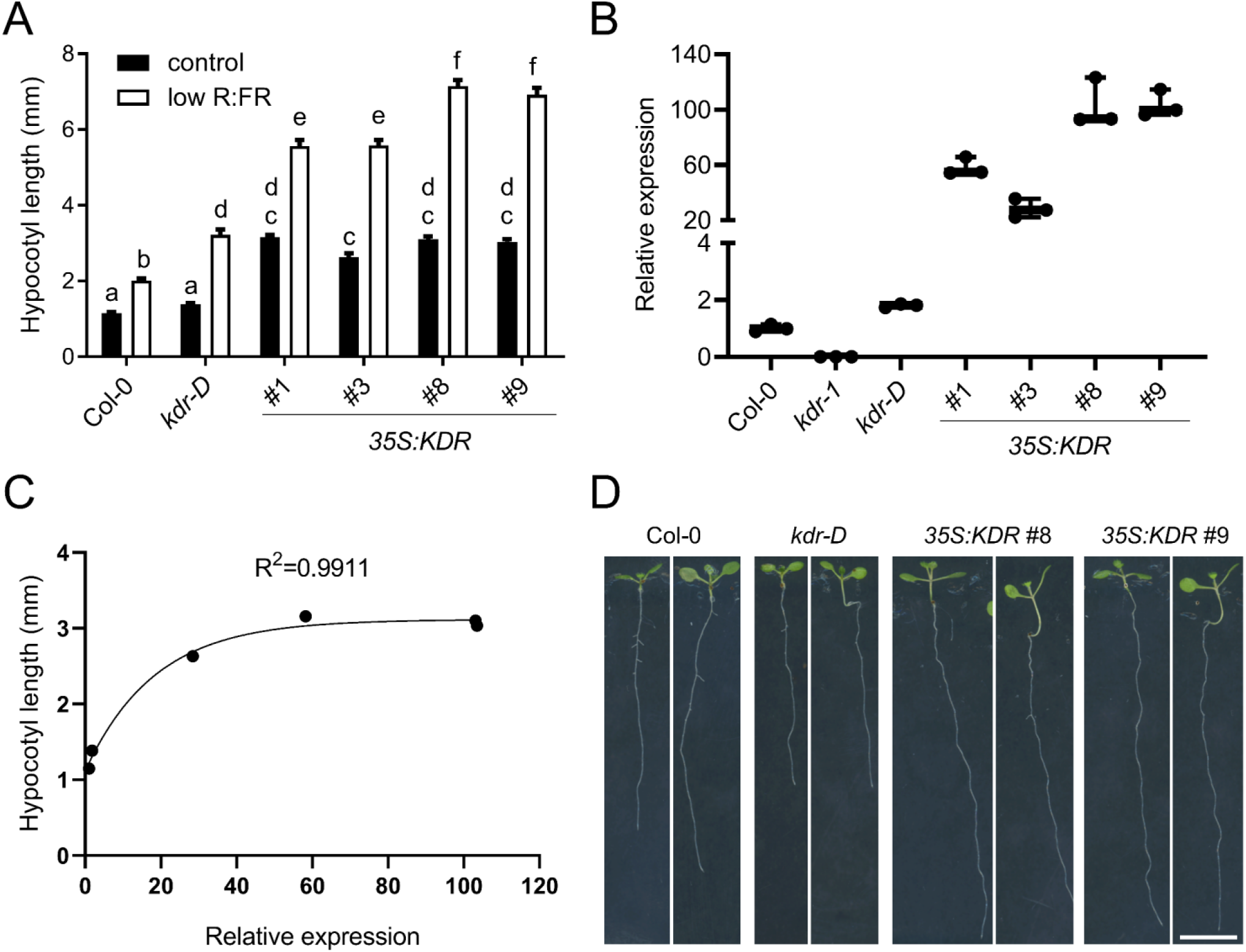
Heterologous overexpression of *KDR* leads to long hypocotyl. (**A**) Hypocotyl length (mm) of seedlings of *A. thaliana* wild type (Col-0), activation-tagged line (*kdr-D*) and independent homozygous transgenic lines overexpressing *KDR* in Col-0 background grown in control light condition (R:FR = 2) or low R:FR (R:FR = 0.2) after 5 days of light treatment. Data represent mean ± SE, n = 38. Different letters indicate statistically significant differences (2-way ANOVA with post-hoc Tukey test, p < 0.05). (**B**) Relative expression level of *KDR* determined by qRT-PCR in wild type Col-0, knockout line (*kdr-1*), activation-tagged line (*kdr-D*) and independent homozygous transgenic lines overexpressing *KDR* in Col-0 background grown in white light. Data represent mean ± SE, n = 3. (**C**) Positive correlation between hypocotyl elongation and expression level of *KDR* measured in Col-0, *kdr-D* and in the transgenic lines overexpressing *KDR* using seedlings grown in control light conditions. The correlation was determined by one-phase association curve fitting, equation: y = y0 + (Plateau – y0) (1 – e^−kx^). Parameters: y0 = 1.115; Plateau = 3.117; k = 0.05431. (**D**) Representative seedlings of hypocotyl length experiment in (**A**), for each genotype the growth is shown in control light (left) and low R:FR (right).

We also verified if petiole elongation in adult plants was affected in our strong overexpression lines. Interestingly, they did not show an increased petiole elongation response to low R:FR treatment (Supplemental Figure 1A). The *KDR* overexpression lines in adult stage are relatively small, but they do seem to have relatively elongated petioles with small leaf laminas (Supplemental Figure 1B), reminiscent of a shade avoidance phenotype. Interestingly, another leaf response, upward movement called hyponasty, was mildly affected in *KDR* overexpression lines (Supplemental Figure 1C). The hyponastic responses of the overexpression lines seemed to be enhanced, especially at an early time point. Finally, *KDR* overexpression lines also exhibit constitutive early bolting and flowering (Figures 3A and 3B and Supplemental Figure 2A), which is another established shade avoidance response (Halliday et al., 1994). The *KDR* overexpressors had very long flowering stems, which at a later life stage started to split open leading to a more bent flowering architecture (Supplemental Figure 2B).

**Figure 3:**
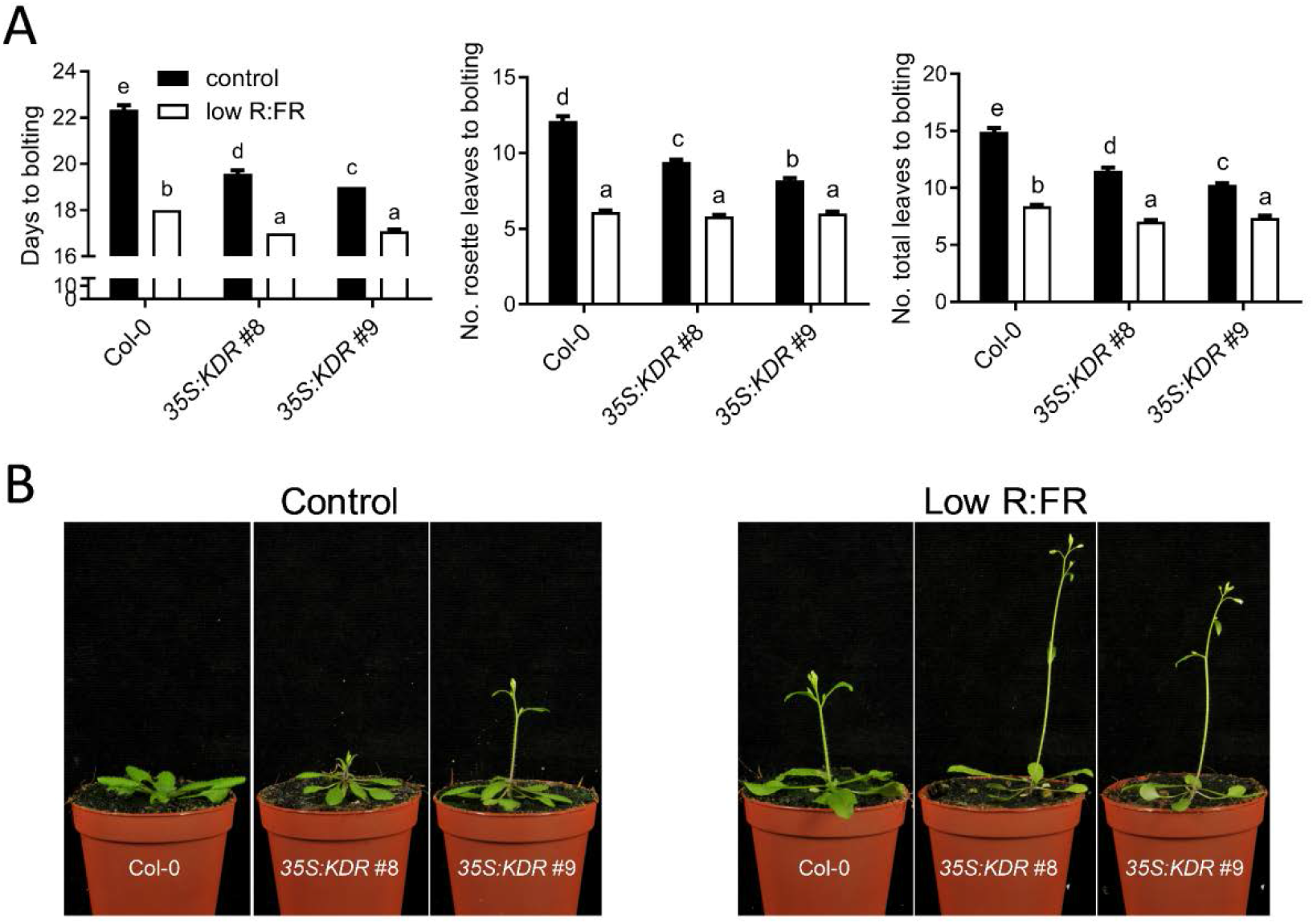
Overexpression of *KDR* leads to a constitutive early flowering. (**A**) Number of days to bolting (left), number of rosette leaves to bolting (middle) and total number of leaves to bolting (right) of *A. thaliana* wild type (Col-0) and two independent transgenic lines overexpressing *KDR* in Col-0 background grown in control light condition (R:FR = 2) or low R:FR (R:FR = 0.2). Data represent mean ± SE, n = 21. Different letters indicate statistically significant differences (2-way ANOVA with post-hoc Tukey test, p < 0.05). (**B**) Representative rosette plants of experiment in (**A**) grown for 20 days in pots containing soil.

### Overexpression of *KDR* affects regulation of PIF targets

The main function of KDR described in literature is its interaction with the negative growth regulator HFR1, which binds to PIFs and therefore interferes with the transcriptional activation of their target genes responsible for the induction of shade avoidance responses. We then verified if some of the well-known PIF targets were transcriptionally regulated in seedling of *A. thaliana* overexpressing *KDR* in control white light conditions in comparison to Col-0. Interestingly, some of the PIF targets were significantly upregulated in the two independent overexpression lines, such as *HFR1, PHYTOCHROME INTERACTING FACTOR 3-LIKE 1* (*PIL1*) and *XYLOGLUCAN ENDOTRANSGLUCOSYLASE/HYDROLASE 17* (*XTH17*), while several other targets, especially the auxin-related genes *YUCCA 8* (*YUC8*), *INDOLE-3-ACETIC ACID INDUCIBLE 29* (*IAA29*) and *IAA19*, were down regulated in the *KDR* overexpression lines compared to Col-0 (Supplemental Figure 3).

### Novel interactors of KDR from Y2H screens

We performed a Y2H screen where the coding sequence (CDS) of *KDR* was cloned in frame with the GAL4 DNA-binding domain of the bait vector and screened against a prey cDNA library of *A. thaliana* cloned with the GAL4 activation domain. The identity of the found interactors are displayed in Table 1, including the frequency with which the interactors were found and the strength of their interaction. Among the proteins identified, we focused on PAR1 and PAR2, two known PIF-interacting proteins. We confirmed the interactions by cloning the full-length CDSs of these proteins (rather than the truncated versions from the library) from *A. thaliana* cDNA into the prey vector and retransformed in yeast to perform a protein-protein interaction assay. In this direct Y2H assay we also tested the previously published interaction of KDR with HFR1 (Hyun and Lee, 2006; Hong et al., 2013), but could not confirm this interaction (Figure 4). Also, when swapping the bait-prey configuration no interaction was found between HFR1 and KDR (Supplemental Figure 4A). Finally, changing vectors to those used in Hyun and Lee, (2006); Hong et al., (2013) (pGBKT7 for the bait and pGADT7 for the prey) would again not confirm the interaction (Supplemental Figure 4B). As positive controls for Y2H assays, we did confirm interactions of HFR1 with PIF4 and PIF5 [previously published in Hornitschek et al. (2009)], and found that HFR1 can also interact with PIF7 (Figure 5A). We also found that KDR does not directly interact with PIFs in yeast, while HFR1 and PAR1 do (Figure 5A). Lastly, we confirmed that PIF7 can interact with itself and other PIFs (Figure 5B), which is consistent with the notion that PIFs form hetero- and homodimers to bind DNA regions and activate the expression of target genes (Bu et al., 2011; Leivar et al., 2008).

**Table 1:**
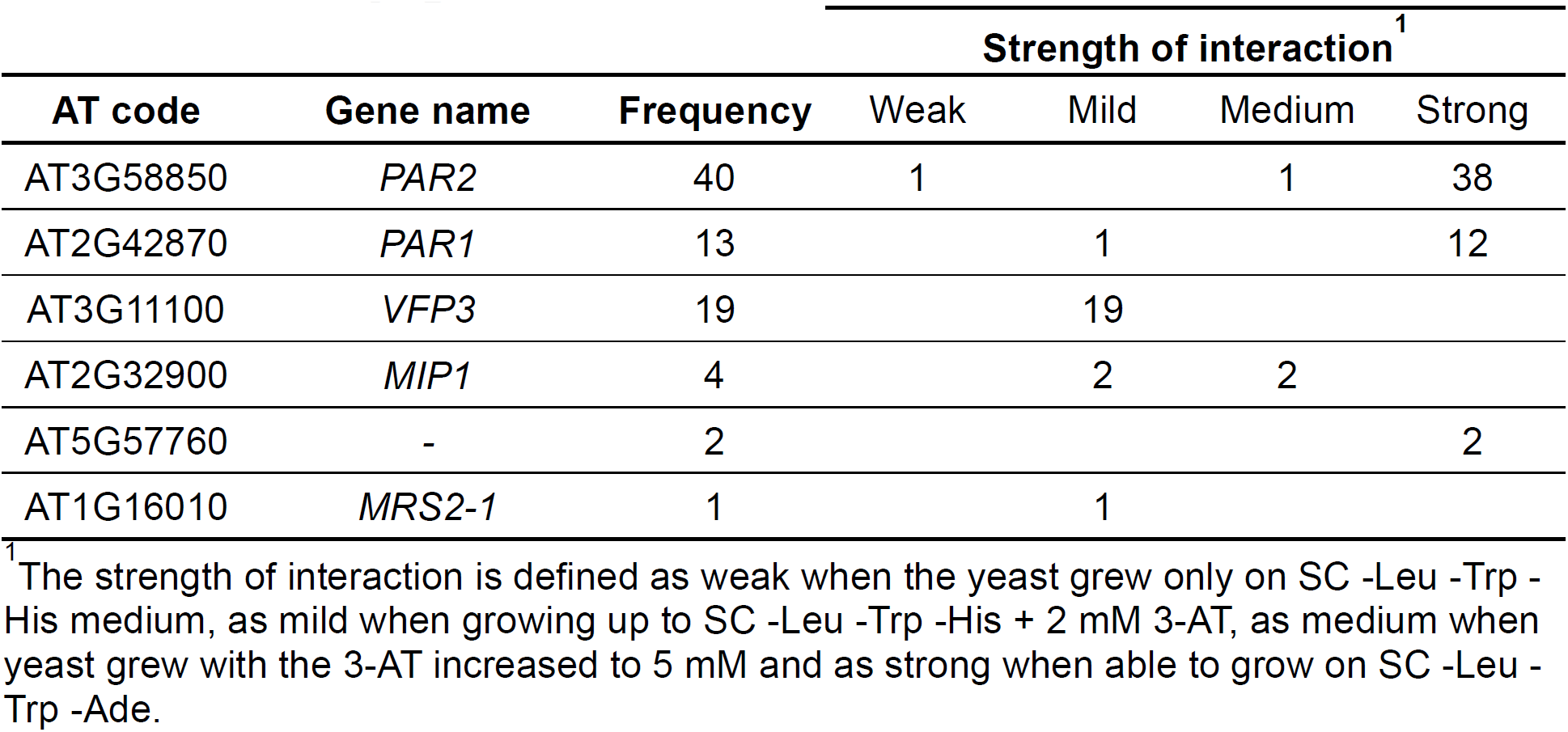
List of candidate interactors identified through a Y2H screening of an *A. thaliana* cDNA library against KDR.

| AT code | Gene name | Frequency | Strength of interaction<sup>1</sup> |  |  |  |
| --- | --- | --- | --- | --- | --- | --- |
|  |  |  | Weak | Mild | Medium | Strong |
| AT3G58850 | *PAR2* | 40 | 1 |  | 1 | 38 |
| AT2G42870 | *PAR1* | 13 |  | 1 |  | 12 |
| AT3G11100 | *VFP3* | 19 |  | 19 |  |  |
| AT2G32900 | *MIP1* | 4 |  | 2 | 2 |  |
| AT5G57760 | - | 2 |  |  |  | 2 |
| AT1G16010 | *MRS2-1* | 1 |  | 1 |  |  |

<sup>1</sup> The strength of interaction is defined as weak when the yeast grew only on SC -Leu -Trp -His medium, as mild when growing up to SC -Leu -Trp -His + 2 mM 3-AT, as medium when yeast grew with the 3-AT increased to 5 mM and as strong when able to grow on SC -Leu -Trp -Ade.

**Figure 4:**
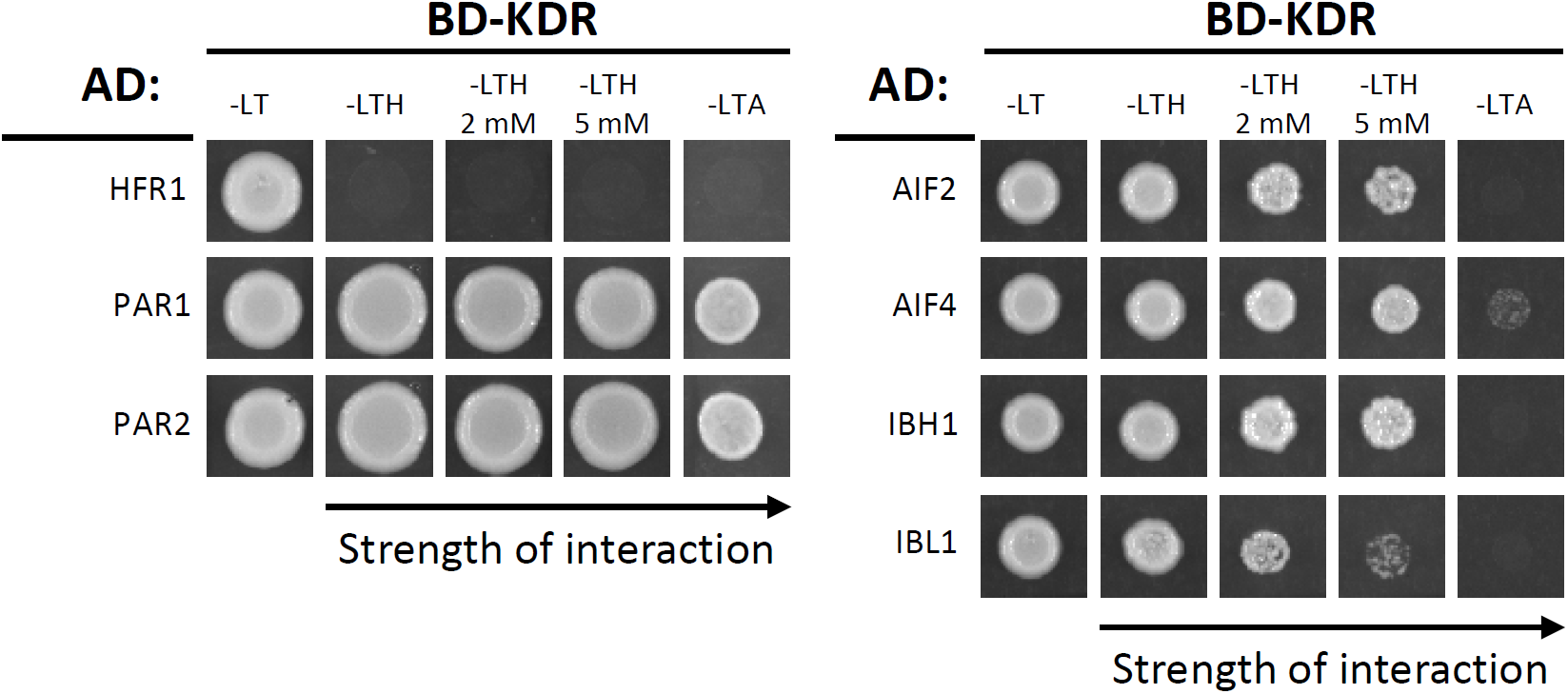
Y2H protein-protein interaction assays confirm the interactions found with screening of two prey libraries of *A. thaliana* against the bait KDR. In the GAL4 Y2H assay, the GAL4-DNA binding domain (BD) fused to KDR was coexpressed with the GAL4-activation domain (AD) fused to the full length CDSs of *HFR1, PAR1, PAR2* (left) and *AIF2, AIF4, IBH1, IBL1* (right). Yeast cells coexpressing the indicated combinations of constructs were grown on nonselective (-LT) or selective (-LTH + 2 or 5 mM 3-AT and – LTA) media. The strength of interaction is shown by the capability of the yeast to grow on stronger selection media, as indicated by the arrow. L: leucine; T: tryptophan; H: histidine; A: adenine; 3-AT: 3-amino-1, 2, 4-triazole.

**Figure 5:**
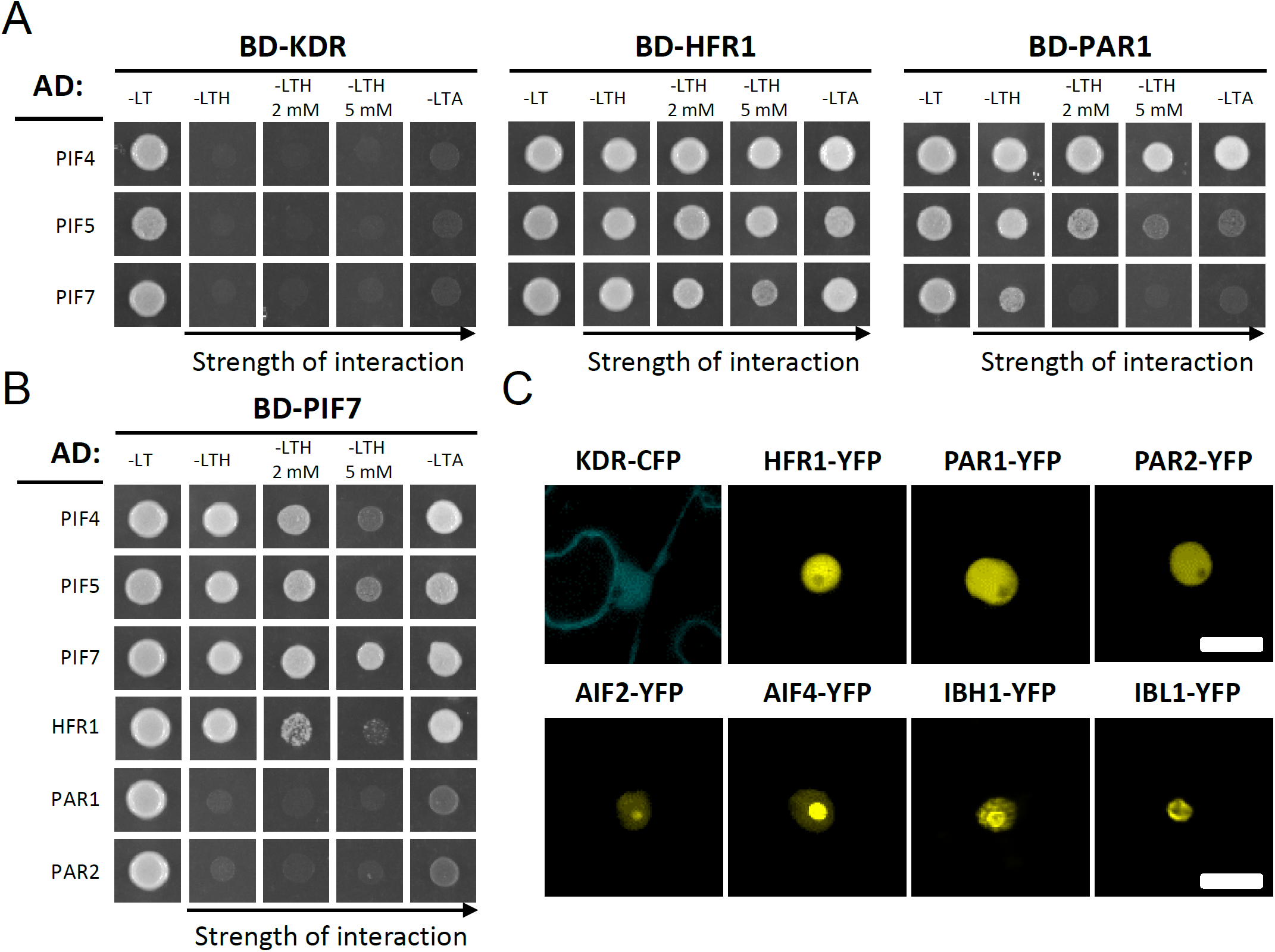
Y2H protein-protein interaction studies and subcellular localization of KDR and its targets *in planta*. (**A**) In the GAL4 Y2H assay, the GAL4 DNA-binding domain (BD) fused to KDR (left), HFR1 (middle) and PAR1 (right) was coexpressed with the GAL4-activation domain (AD) fused to PIF4, PIF5 and PIF7. The mating of the yeast was confirmed through the growth on non-selective medium (–LT). The assay showed that while the bait KDR does not interact with the preys PIF4, PIF5 and PIF7, the bait HFR1 does interact with PIF4, PIF5 and PIF7. PAR1 interacts most strongly with PIF4 and PIF5. (**B**) PIF7 can interact with other PIFs in yeast. When used as bait PIF7 interacts with the prey HFR1 but not with PAR1 and PAR2. (**C**) KDR fused to CFP and the interactors HFR1, PAR1, PAR2, AIF2, AIF4, IBH1 and IBL1 fused to YFP were transiently expressed in epidermal leaf cells of *N. benthamiana* using *A. tumefaciens*. Images were taken 2 days after agroinfiltration. Scale bars indicate 20 µm. L: leucine; T: tryptophan; H: histidine; A: adenine; 3-AT: 3-amino-1, 2, 4-triazole. CFP: cyan fluorescent protein; YFP: yellow fluorescent protein.

In order to maximize the number of relevant KDR interactors found, we performed a second Y2H screen using a completely different library consisting of only TFs of *A. thaliana* cloned in full-length sequence (Pruneda-Paz et al., 2014). Ten putative interactors were discovered and their identity was verified by sequencing (Table 2). We narrowed the selection for further studies to four (b)HLH candidates ATBS1 (ACTIVATION-TAGGED BRI1 SUPPRESSOR 1)-INTERACTING FACTOR 2 (AIF2), AIF4, ILI1 BINDING BHLH 1 (IBH1) and IBH1-LIKE 1 (IBL1), since these proteins had previously been linked to growth regulation in association with some regulators of the SAS, but had not been implemented in shade avoidance control before. The strength of interaction was verified by performing a Y2H direct interaction assay. All four candidates were able to grow at least up to the medium lacking histidine (His) and supplemented with 5 mM 3-amino-1, 2, 4-triazole (3-AT), meaning that the interactions were rather strong in yeast (Figure 4).

**Table 2:** List of candidate interactors identified with a screen of a TF library of *A. thaliana* against KDR.

| AT code | Type | Gene name | Involved in |
| --- | --- | --- | --- |
| AT4G02590 | bHLH | *UNE12* | Fertilization |
| AT2G31210 | bHLH | *bHLH091* | Developing Arabidopsis anther |
| AT2G28160 | bHLH | *FIT* | Regulation of iron uptake responses |
| AT1G64625 | bHLH | *LHL3* | Controlling male meiotic entry |
| AT2G31280 | bHLH155-like protein | *LHL2* | Regulation of early xylem development downstream of auxin |
| AT3G11100 | trihelix | *VFP3* | Interaction with agrobacterium virulence protein VirF |
| AT3G06590 | bHLH | *AIF2* | Negative regulation of cell elongation |
| AT1G09250 | bHLH | *AIF4* |  |
| AT2G43060 | bHLH | *IBH1* | Repressing the expression of PIF4 |
| AT4G30410 | bHLH | *IBL1* |  |

### KDR is localized mainly in the nucleus when coexpressed with strong interactors

Next, we investigated the subcellular localization of KDR and its interactors identified with the Y2H screen. KDR was fused in frame to the N-terminal part of a cyan fluorescent protein (CFP), while the interactors were fused to the yellow fluorescent protein (YFP). Transient expression in *Nicotiana benthamiana* leaves was carried out and revealed that KDR was localized in both the cytoplasm and the nucleus, as previously published by Hong et al. (2013) (Figure 5C). However, the localization of the interactors appeared exclusively in subcellular compartments of the nucleus. In detail, HFR1, PAR1 and PAR2 were detected in the nucleoplasm, while AIF2, AIF4, IBH1 and IBL1 appeared to be localized in the nucleus but with a very pronounced signal in the nucleolus (Figure 5C). When KDR was transiently coexpressed with the putative strong interactors, KDR localized exclusively to the nucleus (Figure 6), indicating that the interaction draws KDR to the nucleus. When coexpressing KDR with HFR1, there was still substantial KDR abundance in the cytoplasm, similar to when KDR was transiently expressed alone, consistent with the lack of interaction found above.

**Figure 6:**
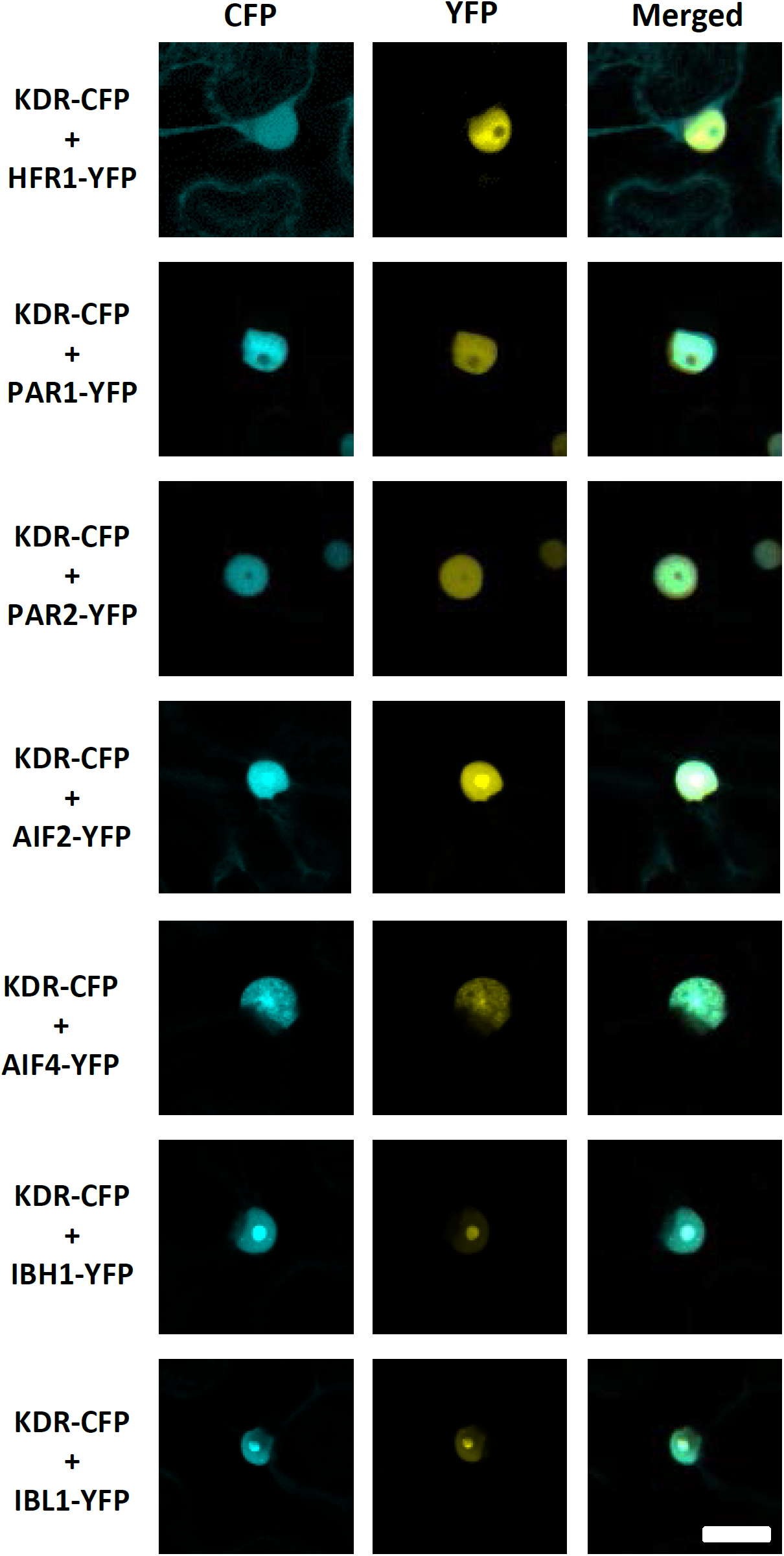
Colocalization of KDR and its interactors *in planta*. KDR and its interactors were transiently coexpressed in epidermal leaf cells of *N. benthamiana*. KDR was fused to CFP, while HFR1, PAR1, PAR2, AIF2, AIF4, IBH1 and IBL1 to YFP. Images were taken 2 days after infiltration with *A. tumefaciens*. The scale bar indicates 20 µm. CFP: cyan fluorescent protein; YFP: yellow fluorescent protein.

### *In planta* BiFC experiments confirm interactions in nuclear compartments

To further verify the interactions of KDR identified with the Y2H assays, we examined whether they were also occurring *in planta*. We performed a BiFC assay where the two parts of the split Venus fluorescent protein (YN or YC) were C- or N-terminally tagged to the proteins of interest and coexpressed in *N. benthamiana* leaves (Figure 7). We detected the reconstituted YFP signal in the nucleus in all the different samples, apart from the interaction with HFR1, and some differences were noticed when KDR was found to interact with the different candidates. The reconstituted YFP signal was observed in different nuclear compartments, resembling the localization of the targets alone (Figure 7). The interactions between KDR and PAR1 and PAR2 were observed in the nucleoplasm while the interactions with AIF2, AIF4, IBH1 and IBL1 were found in the nucleus with the strongest signal in the nucleolus. Together, the Y2H and the BiFC data indicate that KDR can truly interact with all the identified targets and that their interaction seems to trigger its translocation primarily to sub-nuclear complexes, while no interaction with HFR1 could be confirmed.

**Figure 7:**
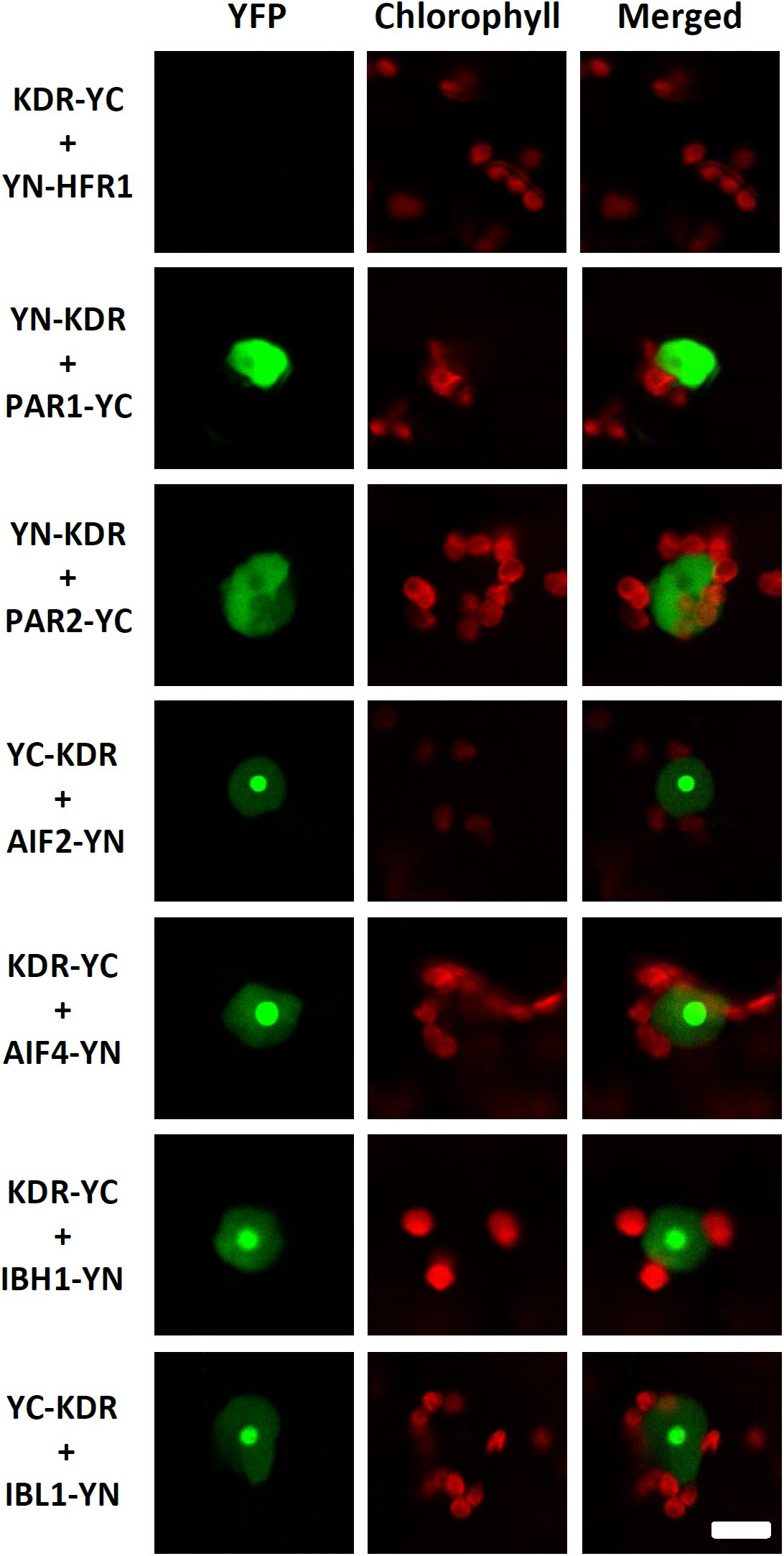
BiFC experiments confirms the interactions found with the Y2H assay. BiFC experiments performed by *A. tumefaciens* transient transformation of *N. benthamiana* leaf epidermis. The interaction of KDR with PAR1, PAR2, AIF2, AIF4, IBH1 and IBL1 was visualized as the reconstituted YFP signal in different nucleus compartments based on the type of interaction. No interaction was found between HFR1 and KDR. The autofluorescence of the chloroplasts is shown in red and the BiFC signal of Venus (YFP) in green. Images were taken 2 days after agroinfiltration. The scale bar represents 10 μm.

### Functional involvement of novel KDR interactors in shade avoidance

HFR1, PAR1 and PAR2 were already associated with shade responses and identified as negative regulators of SAS. Somewhat analogous to KDR, they are transcriptional cofactors, which means they regulate transcription without physically binding DNA but by interacting with other proteins through the HLH domain (Hornitschek et al., 2012; Galstyan et al., 2012, 2011; Roig-Villanova et al., 2007). Also AIF2, AIF4, IBH1 and IBL1 were described as non-DNA-binding (b)HLH proteins (Wang et al., 2009; Ikeda et al., 2012; Zhiponova et al., 2014) but, while their role is mainly related to elongation growth, nothing is known so far about shade avoidance in mutants for these genes. We first confirmed that in our conditions *HFR1, PAR1* and *PAR2* were also upregulated following a low R:FR treatment in seedlings of Col-0 (Figure 8A). Since *AIF2, AIF4, IBH1* and *IBL1* were never associated with shade responses, we also verified if their expression level was differentially regulated upon exposure to low R:FR and we found that this was indeed the case (Figure 8A).

**Figure 8:**
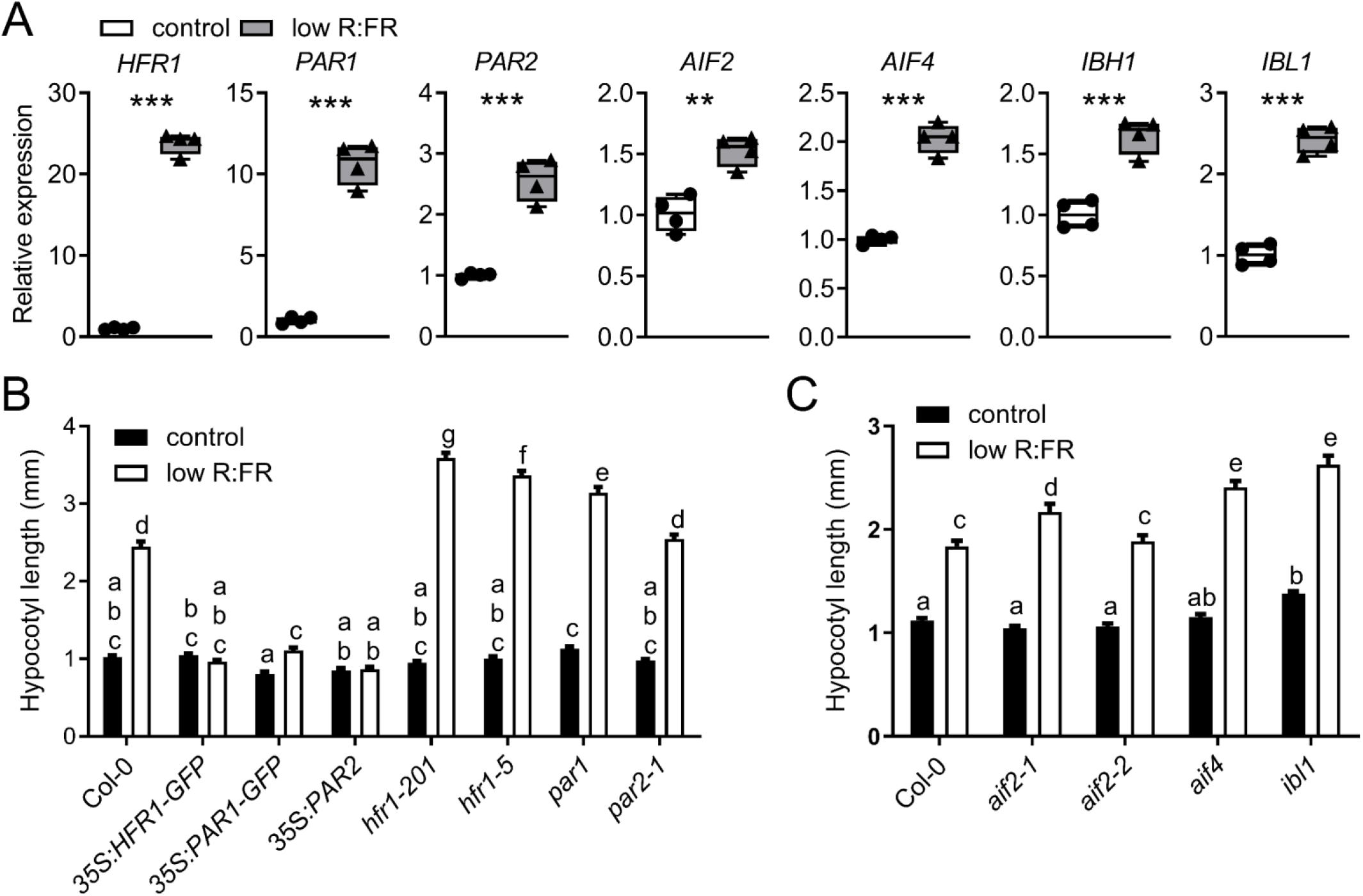
KDR interactors are differentially expressed and their misexpression affects phenotypic responses to low R:FR. (**A**) Relative expression of *HFR1, PAR1, PAR2, AIF2, AIF4, IBH1* and *IBL1* in control white light or low R:FR light for 90 minutes in wild type Col-0 shoots. Data represent mean ± SE, n = 4. Statistically significant difference indicated by *** for p < 0.001 and ** for p < 0.01, Student’s t-test. (**B** and **C**) Hypocotyl length (mm) of seedlings of *A. thaliana* wild type (Col-0), overexpressing lines (*35S:HFR1-GFP, 35S:PAR1-GFP* and *35S:PAR2*) and mutants (*hfr1-201, hfr1-5, par1, par2-1, aif2-1, aif2-2, aif4* and *ibl1*) grown in control light condition (R:FR = 2) or low R:FR (R:FR = 0.2) after 5 days of light treatment. Data represent mean ± SE, n = 38. Different letters indicate statistically significant differences (2-way ANOVA with post-hoc Tukey test, p < 0.05).

Next, we studied the response to low R:FR conditions of different mutant and overexpression lines of these bHLH genes relative to Col-0 wild type (Figures 8B and 8C). Figure 8B shows that low R:FR-induced hypocotyl elongation is increased in the *hfr1-201, hfr1-5* and *par1* mutants. Overexpressing any of these genes was sufficient to severely block the response to low R:FR. These phenotypes are entirely consistent with the roles of these proteins as negative SAS regulators. Instead, for the less known genes *AIF2, AIF4, IBH1* and *IBL1* we studied the available T-DNA insertion lines, but unfortunately in the SALK lines for *IBH1* we could not detect the T-DNA insertion and were therefore discarded. In low R:FR conditions, all the lines except for *aif2-2* showed a moderately enhanced hypocotyl elongation response compared to Col-0 wild type (Figure 8C). In the case of *ibl1*, a statistically significant difference, although minimal, was already seen in control conditions when compared to the wild type. The relatively mild phenotypes, albeit reproducible and statistically significant, may hint at genetic redundancy between the different KDR targets. Higher order combinations of these mutants, as well as overexpression lines for these genes, would likely help understand the impact of these novel shade avoidance components in more detail.

### Genetic interaction between KDR and downstream shade avoidance regulators

We hypothesized that KDR would act to sequester negative regulators, such as PAR2, by direct interaction. We verified this hypothesis by crossing a *PAR2* overexpression line with a *KDR* overexpression line. The high KDR abundance should sequester PAR2 and restore elongation growth in the *PAR2* overexpressor (Figures 9A and 9B). Indeed, we showed that *KDR* overexpression fully rescues hypocotyl elongation and response to low R:FR in the *PAR2* overexpression line. Interestingly, *KDR* overexpression could not rescue low R:FR-induced hypocotyl elongation in an *HFR1* overexpression line. These data are fully consistent with the observations that KDR interacts with PAR2, but does not seem to interact with HFR1. Comparable results were also found when *PAR2* and *HFR1* overexpressor lines were crossed with the mild overexpression line *kdr-D*. Also this time the *HFR1* overexpressor phenotype could not be rescued, while the elongation of the hypocotyl of *PAR2* overexpressor was comparable to that of *kdr-D*. A closer look at this cross shows that the appearance of the cotyledons and their “petioles” are similar to those exhibited by *35S:PAR2* overexpression line (Supplemental Figure 5).

**Figure 9:**
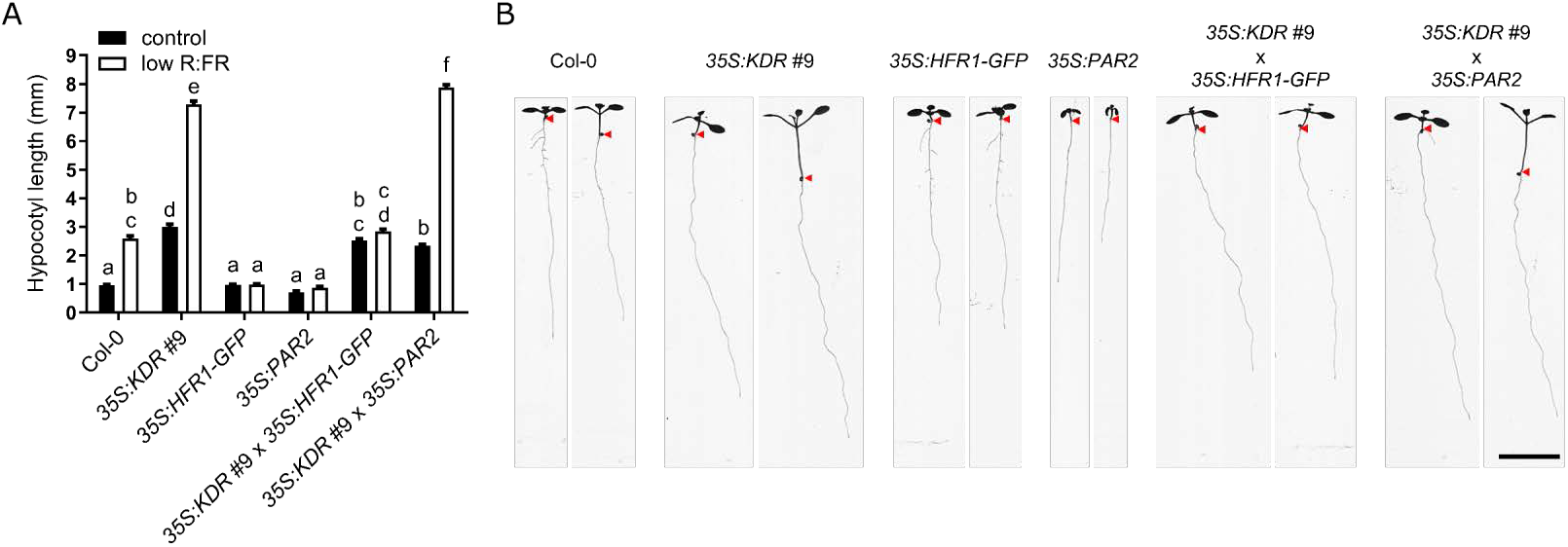
Overexpression of *KDR* rescues hypocotyl length in response to low R:FR in *PAR2* but not *HFR1* overexpressor. (**A**) Hypocotyl length (mm) of seedlings of *A. thaliana* wild type (Col-0), *35S:KDR* #9; *35S:HFR1-GFP* and *35S:PAR2* overexpression lines and the double overexpression lines *35S:KDR* #9 × *35S:HFR1-GFP* and *35S:KDR* #9 × *35S:PAR2* grown in control light condition (R:FR = 2) or low R:FR (R:FR = 0.2) after 5 days of light treatment. Data represent mean ± SE, n = 38. Different letters indicate statistically significant differences (2-way ANOVA with post-hoc Tukey test, p < 0.05). (**B**) Representative seedlings as in experiment (**A**), for each genotype the growth is shown in control light (left) and low R:FR (right). The arrows indicate the hypocotyl-root transition. The scale bar represents 1 cm.

Finally, we also generated transgenic lines overexpressing *KDR* in *pif7, pif4 pif5* and *pif4 pif5 pif7* backgrounds by floral dipping these mutants with a *35S:KDR* construct using Agrobacterium-mediated transformation. We then studied their response when exposed to low R:FR using three independent lines for each background, after we verified their expression level (Figure 10 and Supplemental Figure 6). As expected, *pif4 pif5* shows a clear but reduced response to low R:FR, while *pif7* and *pif4 pif5 pif7* lost the hypocotyl response to low R:FR completely. Interestingly, in control white light conditions the overexpression of *KDR* is able to induce a strong elongation in *pif7* and *pif4 pif5* backgrounds, and more mildly when overexpressed in the triple knockout *pif4 pif5 pif7*. When these lines were exposed to low R:FR, *pif4 pif5 35S:KDR* had nearly the same hypocotyl phenotype as the same construct has in wild type background, consistent with the relatively modest role of *pif4 pif5* in low R:FR-induced hypocotyl elongation. Contrary, in the *pif4 pif5 pif7* mutant, *KDR* overexpression could not rescue the hypocotyl elongation response to low R:FR, while in the *pif7* only mildly.

**Figure 10:**
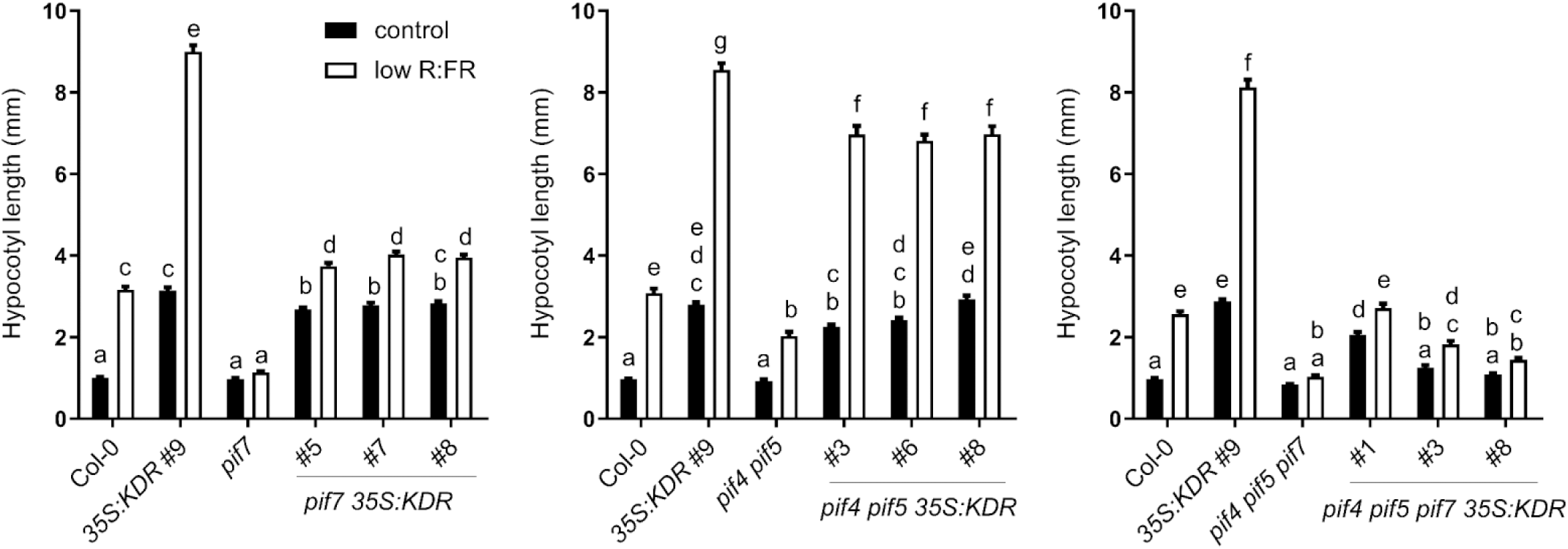
Overexpression of *KDR* in different *pif* knockout lines. Hypocotyl length (mm) of seedlings of *A. thaliana* wild type (Col-0), knockout lines (*pif7, pif4 pif5* and *pif4 pif5 pif7*), *KDR* overexpressing line (*35S:KDR* #9) and independent lines of *pif7, pif4 pif*5 and *pif4 pif5 pif7* overexpressing *KDR* (*pif7 35S:KDR, pif4 pif5 35S:KDR* and *pif4 pif5 pif7 35S:KDR*) in control light condition (R:FR = 2) or low R:FR (R:FR = 0.2) after 5 days of light treatment. Data represent mean ± SE, n = 38. Different letters indicate statistically significant differences (2-way ANOVA with post-hoc Tukey test, p < 0.05).

We conclude that KDR interacts with PARs to regulate hypocotyl elongation in response to low R:FR and this subsequently depends on PIFs, probably because PARs directly interact with PIFs to regulate their activity.

## DISCUSSION

Significant discoveries have been made in the past decades to identify the molecular mechanisms through which plants perceive neighbors through light signals and activate the shade avoidance network. A previous transcriptome analysis on two wild *Geranium* species, one shade tolerant, while the other shade avoiding, identified *KDR* as a molecular component whose expression was correlated with the ability to display elongation responses to shade (Gommers et al., 2017). Here we attempted to unravel its role in SAS and found that overexpression of *KDR* in *A. thaliana* resulted in an enhanced response to simulated shade conditions. Therefore, it was proposed that the interaction of KDR with HFR1 would release PIFs so that they could activate shade avoidance. If this would have been the only mode of action of KDR, then lines overexpressing *KDR* should have a similar phenotype to knockout lines of *hfr1*, which is not the case. Moreover, independent studies on non-model plants identified *KDR* orthologues as strong candidate regulators of contrasting elongation responses and in these species no *HFR1* orthologues could be found (van Veen et al., 2013; Gommers et al., 2017). Therefore, we speculated that other interacting partners might exist.

### KDR is a novel regulator of established shade avoidance components

Results from one of the Y2H screens and further *in planta* confirmations identified PAR1 and its closest homolog PAR2 as interactors of KDR, proteins that were previously associated with SAS (Zhou et al., 2014; Cifuentes-Esquivel et al., 2013; Hao et al., 2012; Galstyan et al., 2012, 2011; Bou-Torrent et al., 2008; Roig-Villanova et al., 2007). However, their interaction with KDR was not previously anticipated and this sheds new light on shade avoidance control. PAR1 and PAR2, as well as KDR and HFR1, are atypical non-DNA-binding (b)HLHs. HFR1 regulates shade avoidance by interacting with several PIFs (PIF1, PIF3, PIF4 and PIF5). This yields non-functional complexes unable to bind DNA and therefore blocks the activation of their targets (Hornitschek et al. 2009; Zhang et al. 2013). We show here that HFR1 can also interact with PIF7 (Figures 5A and 5B). Inactivation of PIF4 also occurs via interaction with PAR1 and PAR2, adding an extra level of regulation of cell elongation and plant development. Furthermore, PAR1 can also interact with BES1-INTERACTING-MYC-LIKE 1 (BIM1) and with the BRASSINOSTEROID-ENHANCED EXPRESSION 1 (BEE1), BEE2 and BEE3, which are positive regulators of SAS, by forming non-functional complexes also in this case (Cifuentes-Esquivel et al., 2013). Finally, overexpression of *PRE1*, another (b)HLH member of the same subgroup as KDR, can suppress the dwarf phenotype of *PAR1* overexpression (Hao et al., 2012). In a comparable way, we found that overexpression of *KDR* can restore the growth defect of *PAR2* overexpression (Figure 9). This finding places KDR in a new third level of SAS regulation, above PAR1 and PAR2, which suppress PIF activity (Figure 11). Our results also cast severe doubts on the suppressing role of KDR on HFR1, at least in the regulation of SAS.

**Figure 11:**
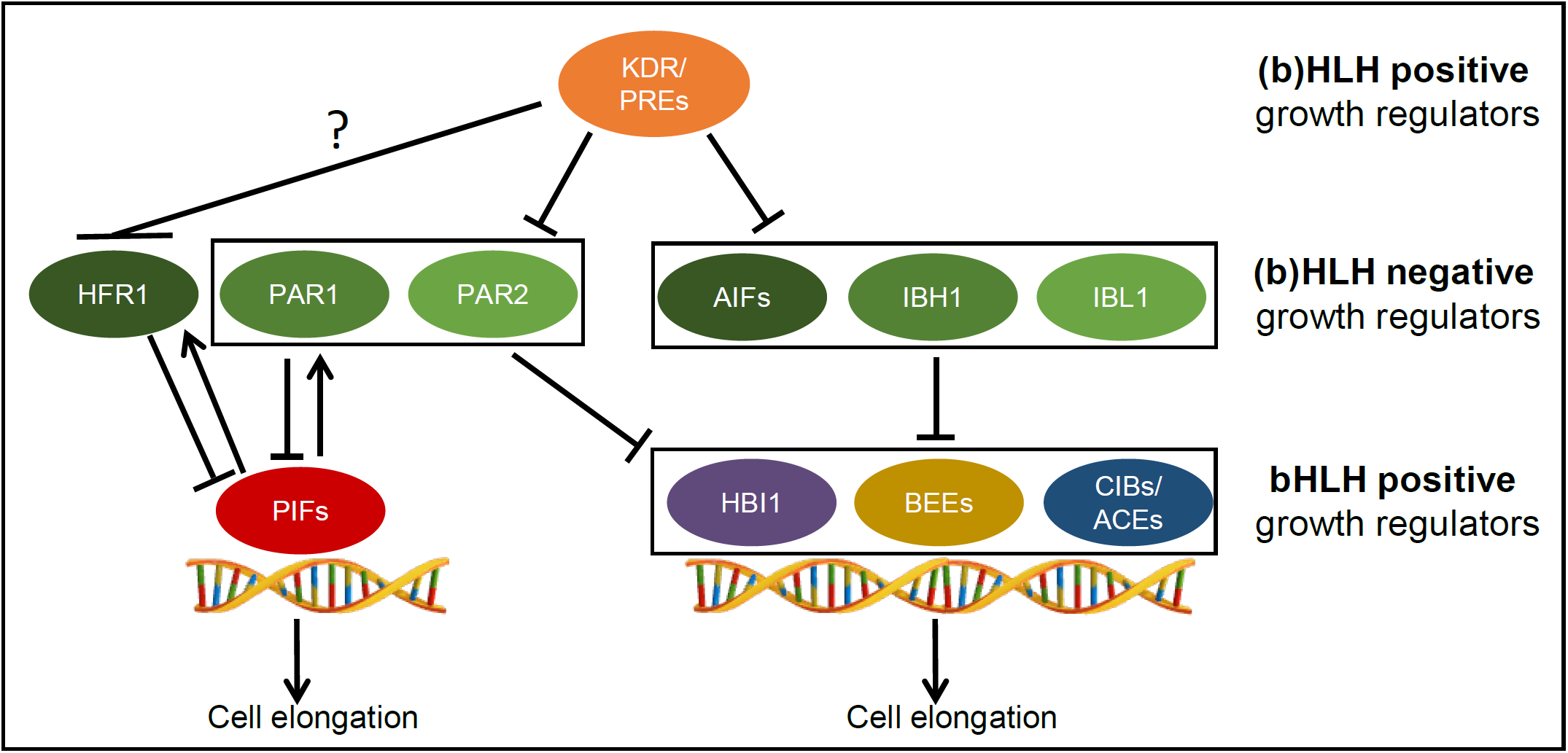
Proposed network regulating cell elongation in low R:FR. The HHbH module is composed by atypical (b)HLH and typical bHLH members, which can be positive or negative growth regulators and they interact in an antagonistic and redundant manner to regulate adequately and rapidly cell elongation.

### KDR interacts with several negative growth regulators

In this study, we show that KDR physically interacts in yeast and *in planta* with a range of negative regulators of cell elongation, *i.e.* AIF2, AIF4, IBH1 and IBL1. None of them had been previously associated with SAS but they all share some similarities. For example, they had been identified already for their interaction with some of the PRE members (Wang et al., 2009; Ikeda et al., 2013; Zhang et al., 2009); they are atypical (b)HLH proteins, unable to bind DNA (Ikeda et al., 2012, 2013) and their overexpression results in a dwarf phenotype (Ikeda et al., 2013; Zhang et al., 2009). Nevertheless, little is still known about these proteins. They have been found to interact with growth-promoting TFs, possibly to suppress the elongation growth (Ikeda et al., 2012; Bai et al., 2012), similar to the mode of action of PARs and HFR1. Furthermore, the dwarf phenotype of *IBH1* overexpression is restored when crossed with an overexpressing line of a *PRE* member (Zhang et al. 2009), reminiscent of the *35S:KDR* #9 × *35S:PAR2* cross shown here (Figure 9).

In conclusion, KDR interacts with and sequesters (b)HLH transcription cofactors, such as PARs, AIFs, IBH1 and IBL1, which are negative growth regulators. PARs, AIFs, IBH1 and IBL1 bind to the bHLH TFs PIFs, ACTIVATOR FOR CELL ELONGATION 1-3 (ACE1-3), CRYPTOCHROME-INTERACTING BASIC-HELIX-LOOP-HELIX 5 (CIB5), HOMOLOG OF BEE2 INTERACTING WITH IBH1 (HBI1) and BEE2, likely inhibiting their DNA-binding activities to promote cell elongation. Upon binding, KDR would then sequester PARs, AIFs, IBH1 and IBL1 so they are kept from binding their bHLH TF targets. This triantagonistic network of bHLHs fine tunes general plant development as well as adaptation to environmental changes, *i.e.* changes in light quality. Although under natural conditions, each of these proteins have subtle impacts on development and plasticity, overexpression of these factors leads to a severe dwarfing phenotype and impaired cell elongation, in agreement with their inhibition of positive growth regulators (Ikeda, Mitsuda, and Ohme-Takagi 2013; Zhang et al. 2009; Zhiponova et al. 2014). On the contrary, ectopic expression of *KDR* leads to a strongly elongated phenotype. To test if the activity of these novel negative growth regulators is really inhibited by interaction with KDR, lines overexpressing *KDR* and the negative growth regulators at the same time could test this hypothesis, similar to our data here for combined *PAR2* and *KDR* overexpression (Figure 9).

### The heterodimerization of KDR leads to its nuclear translocation

The interaction of KDR with the (b)HLH cofactors shown here is also reinforced by its translocation from the cytoplasm to the nucleus when coexpressed with the interacting bHLHs identified in the Y2H (Figure 6). Since bHLH proteins are a family of TFs, they are thought to be localized mostly in the nucleus where they can regulate the transcription of genes. On the other hand, literature presents evidence showing that the translocation from the cytoplasm to the nucleus is an important post-translational regulatory mechanism in response to different stimuli and to different plant developmental stages (McGonigle et al., 1996; Nayar et al., 2014; Cui et al., 2016). A possible explanation for KDR translocation is that it could have a weak nuclear localization signal (NLS), while its interactors could have a strong NLS and therefore are localized only in the nucleus even in absence of KDR. Consequently, the predominantly nuclear localization of KDR relies on partners harboring strong NLS. Indeed, this is another possible mechanism of modulating SAS. In fact, all the strong interactors identified here are upregulated following exposure to low R:FR, suggesting that KDR could change its localization during shade to form heterodimers with the negative growth regulators, inhibiting their function and having as final output the promotion of cell elongation through the release of the other positive growth regulators.

### Several triantagonistic (b)HLH/(b)HLH/bHLH (HHbH) modules control cell elongation in response to multiple stimuli

Taken together, we propose that the response to shade is mediated by the coaction of several modules of heterodimers formed between bHLH proteins with opposite function in the regulation of elongation (Figure 11). Furthermore, we conclude that KDR can regulate a substantial number of shade avoidance regulators, certainly many more than previously anticipated, and this would explain the profound impact of variations in *KDR* expression levels on shade avoidance. Some of them are well known negative regulators of shade avoidance responses, *i.e.* PAR1 and PAR2 (Roig-Villanova et al., 2007; Roig-Villanova et al., 2006; Sessa et al., 2005), while other interactors found, *i.e.* AIF2, AIF4, IBH1 and IBL1, had never been associated with shade avoidance. The expression levels of all these novel KDR interactors were induced upon low R:FR treatment, consistent with their involvement in the regulation of shade avoidance (Figure 8A).

The results described here uncovered new levels of shade avoidance regulation, while at the same showing that HFR1 and KDR likely act independently. The gas-and-brake mechanism of different layers of (b)HLH proteins described here gives tremendous opportunity to fine-tuning shade avoidance expression. This could be instrumental to enhancing low R:FR response by simultaneous blue light deprivation (de Wit et al., 2016) or suppressing it during abiotic stress (Hayes et al., 2019). Unraveling the exact roles of these novel bHLH interactions through different life stages of the plant, under different light conditions and in other stress response pathways would enable an integrative understanding of plant shade avoidance.

## METHODS

### Plant material, growth conditions and measurements

*A. thaliana* lines are all in Col-0 background. The following published lines were used: the activation-tagged line *kdr-D* and the knockout *kdr-1* (SALK_048383C) (Hyun and Lee, 2006; Gommers et al., 2017), *35S:HFR1-GFP* (also called *G-BH* 03) (Galstyan et al., 2011), *35S:PAR1-GFP, hfr1-201* (Zhou et al., 2014), *35S:PAR2, par2-1* (Roig-Villanova et al., 2007), *hfr1-5* (Sessa et al. 2005), *par1* (SAIL_668_E10, Khanna et al. 2006), *pif7-1* (Leivar et al., 2008), *pif4-101 pif5-1* (Lorrain et al., 2008), *pif4-101 pif5-1 pif7-1* (de Wit et al., 2015). Instead, seeds of *aif2-1* (SALK_ 011076C), *aif2-2* (SALK_ 061834), *aif4* (GK-428G06) and *ibl1* (SALK_119457C, Zhiponova et al., 2014) were obtained from The Nottingham Arabidopsis Stock Centre (NASC, UK) and genotyped using the primers listed in Supplemental Table 1. *35S:HFR1-GFP × kdr-D* and *35:PAR2 × kdr-D* were created by crossing the respective genotypes, described above, and experiments were performed using lines homozygous for both inserts.

The plants were cold stratified for 4 days on soil:perlite mix 1:2 (Primasta BV, Asten, The Netherlands) supplemented with nutrients (2.6 mM KNO_3_, 2 mM Ca[NO_3_]_2_, 0.6 mM KH_2_PO_4_, 0.9 mM MgSO_4_, 6.6 µM MnSO_4_, 2.8 µM ZnSO_4_, 0.5 µM CuSO_4_, 66 µM H_3_BO_3_, 0.8 µM Na_2_MoO_4_ and 134 mM Fe-EDTA; pH 5.8) and then moved to growth chamber under short-day conditions (8 h light, 16 h dark; 20°C; 70% humidity; PAR ∼ 140 µmol m^−2^ s^−1^) for 11 days to allow germination. The plants were then transplanted and left to grow for 3 weeks until the beginning of the experiment. In the case of long-day conditions (16 h light, 8 h dark), the plants were grown for 7 days before being transplanted and then left to grow for an extra week before the light treatment started. Petiole length of the third youngest leaf was measured with a digital caliper before and after the treatment and the difference calculated. Pictures of the same plants were taken before (t = 0) and after the treatment (t = 8 h and t = 24 h), and the Δ in petiole angle was measured with ImageJ to determine the hyponastic response. For bolting and flowering experiments, a single seed per pot was sowed, cold stratified for 4 days and then moved to the growth chamber under long-day conditions (16 h light, 8 h dark) for 4 days to allow germination before the light treatment started. The number of days to bolting and to flowering was calculated starting from the first day the pots were moved to the growth chamber until the flowering bolt appeared and the first flower opened, respectively. The number of rosette leaves and the total number of leaves (rosette leaves plus cauline leaves) were also measured at the moment of bolting and flowering.

For experiments on *A. thaliana* seedlings, the seeds were gas sterilized with chloride for 2 h, sown on sterile square Petri dishes (120 × 120 × 17 mm, Greiner Bio One) containing ½ MS (Duchefa Biochemie), 1 g/L MES hydrate (Sigma-Aldrich), 8 g/L Plant agar (Duchefa Biochemie) at pH 5.8 and cold stratified for 4 days. The plates were placed in the light for 2 h followed by one day of darkness and then back to the light for 2 days, after which the light treatment started. The plates were finally scanned, and the hypocotyls were measured with ImageJ.

*N. benthamiana* seeds were germinated for 7 days (16 h light, 8 h dark; 20°C; 70% humidity; PAR ∼140 µmol m^−2^ s^−1^) in 9 × 9 × 9.5 cm pots containing soil:perlite mix (1:2) (Primasta BV, Asten, The Netherlands). The seedlings were then transplanted to 7 × 7 × 8 cm pots and grown for 4-5 more weeks before agroinfiltration experiments were started.

### Light treatment

For simulated shade conditions, FR LEDs (730 nm, Philips GreenPower) were used to decrease the R:FR ratio from 2 (control conditions) to 0.2 (shade conditions) without changing the PAR (140 µmol m^−2^ s^−1^, Philips HPI 400 W). The light treatment lasted 5 days when using *A. thaliana* seedlings, and 24 h for adult *A. thaliana* plants. The light spectra of the different light conditions are shown in Supplemental Figure 7.

### RNA extraction and qRT-PCR

Whole seedling shoots of 4-day-old seedlings growing on plates (Figures 1A, 8A and Supplemental Figure 3) and whole seedling shoot of 8-day-old seedling (Figure 2C and Supplemental Figure 6) were used to extract RNA using the RNeasy® Mini kit (Qiagen, Hilden, Germany) followed by DNase I (Qiagen, Hilden, Germany) treatment. cDNA synthesis was performed using SuperScript™ III RNase H^−^ Reverse Transcriptase (Thermo Fisher Scientific) together with random primers (Invitrogen, Waltham, USA). qRT-PCR reactions were conducted in Viia7 Real-Time PCR (Thermo Fisher Scientific) using the SYBR™ Green Supermix (Bio-Rad, Hercules, USA). Two technical replicates of three or four biological samples were used to calculate the average gene expression level normalized to the housekeeping genes *AT4G26410* and *AT5G25760* and relative to the expression level of wild type Col-0 control condition. The primers used for qRT-PCR are listed in Supplemental Table 2.

### Gene cloning and plant transformation

The cDNA used to clone *KDR* CDS of *A. thaliana* Col-0 was synthesized from RNA derived from leaf. The CDS was amplified using the Phusion DNA polymerase (Thermo Fisher Scientific) with the *att*B primers listed in Supplemental Table 3 and cloned into the Gateway vector pDONR207 (Thermo Fisher Scientific) using the Gateway BP clonase II enzyme mix (Thermo Fisher Scientific). The reaction was used to transform competent cells of *Escherichia coli* DH5α. Colonies growing on the selective medium containing the antibiotic gentamycin (20 µg/ml) were checked by colony PCR and the plasmid DNA extracted using the QIAprep® Spin Miniprep kit (Qiagen). A restriction reaction was performed on the extracted plasmid DNA and the positive samples were sequenced. The entry vector with the right sequence was recombined into the Gateway destination vector pFAST-G02 (Shimada et al., 2010) using the Gateway LR clonase II enzyme mix (Thermo Fisher Scientific). The reaction was then used to transform *E. coli* DH5α competent cells. The colonies growing on the antibiotics streptomycin and spectinomycin (100 µg/ml each) were checked by colony PCR. The plasmid DNA was extracted and double checked with restriction reaction. Less than 10 ng of construct was used to transform competent cells of *Agrobacterium tumefaciens* AGL-1 that were able to grow on the antibiotic rifampicin (20 µg/ml). Positive colonies growing on the selective antibiotics were confirmed by colony PCR and then used to transform flowering plants of *A. thaliana* Col-0, *pif4, pif4 pif5* and *pif4 pif5 pif7* following the protocol of Zhang et al. (2006). Successfully transformed T_1_ seeds were selected through the GFP signal in dry seeds. T_2_ lines were selected for single insertion of the transgene using the selectable marker *bar*, which confers resistance to the herbicide Basta (25 µg/ml) (DL-Phosphinothricin, Duchefa Biochemie). Finally, T_3_ seeds were screened for homozygosity using the GFP signal and the insertion of the transgene was confirmed by PCR reaction performed on genomic DNA (gDNA) extracted from homozygous plants using the primers listed in Supplemental Table 4. Experiments were performed using T_3_ or T_4_ seeds.

### TAIL-PCR

Leaf material was used to extract gDNA from transgenic lines *35S:KDR* #1, 3, 8 and 9, created using the pFAST-G02 vector, as described above. The gDNA was used to perform a TAIL-PCR reaction as described in Liu et al. (1995) with minor modifications, using arbitrary degenerate (AD), T-DNA left border (LB) end primers (Supplemental Table 5) and DreamTaq DNA polymerase (Thermo Fisher Scientific). The cycle settings used for the TAIL-PCR reactions were adjusted based on the characteristics of the polymerase and primers used and listed in Supplemental Table 6. Purified fragments obtained in the second or third TAIL-PCR reactions were sequenced (Macrogen Europe, Amsterdam) and analyzed through a BLAST search (NCBI, www.ncbi.nlm.nih.gov) to identify the flanking sequences. A schematic representation of the insertion sites is shown in Supplemental Figure 8.

### Gene cloning for Y2H interactions

The procedure for cloning *KDR* CDS was as described before but with the use of the primers listed in Supplemental Table 7 and cloned into the Gateway vector pDONR221 (Thermo Fisher Scientific). The competent cells transformed with the entry vectors were selected for growth on the antibiotic kanamycin (50 µg/ml). The CDSs of the genes *HFR1, PIF4, PIF5* and *PIF7* cloned into the Gateway vector pENTR/D-TOPO and of *PAR1* and *PAR2* cloned into the Gateway vector pENTR223 were obtained from ABRC and sequence-validated. The entry vectors containing the CDSs of *KDR, HFR1, PAR1, PAR2, PIF4, PIF5* and *PIF7* were recombined into the Gateway destination vector pDEST32 (gentamycin 20 µg/ml) while *KDR, HFR1, PAR1, PAR2, PIF4, PIF5* and *PIF7* were also recombined in the Gateway destination vector pDEST22 (carbenicillin 50 µg/ml). The pDEST22 vectors harboring the CDS of the genes *AIF2, AIF4, IBH1* and *IBL1* were obtained from the yeast TFs library described by Pruneda-Paz et al. (2014). The yeast colonies were grown on 5 ml liquid synthetic complete (SC) medium (Formedium, Hunstanton, UK) lacking the selective amino acid (AA) tryptophan (Trp) for one day at 30°C in shaking conditions. The plasmid DNA was then extracted using the QIAprep® Spin Miniprep kit (Qiagen). Finally, the entry vectors containing *KDR* and *HFR1* were both cloned also into the Gateway destination vectors pGBKT7 (kanamycin 50 µg/ml) and pGADT7 (carbenicillin 50 µg/ml).

### Yeast prey plasmid cDNA library of *A. thaliana*

The yeast prey plasmid cDNA library of *A. thaliana* was kindly provided by Prof. Dr. Guido van den Ackerveken (Utrecht University, The Netherlands) and created using Invitrogen Custom Services (Invitrogen, Carlsbad, CA). RNA was extracted from 15-day-old *A. thaliana* seedlings subjected to five conditions: uninfected, infected with a compatible strain of *Hyaloperonospora Arabidopsidis* (*Hpa*), infected with incompatible strain of *Hpa*, sprayed with benzothiadiazole (BTH) or infiltrated with NIN-like proteins (NLPs). This variety of optimal and stress conditions likely yielded a very broad library of different transcripts. Synthesis of cDNA was performed on the RNA extracted, cloned into the Gateway donor vector pENTR222 and then recombined into the Y2H Gateway destination vector pDEST22 to generate a GAL4 activation domain (AD) fused to the N-terminal part of *A. thaliana* proteins. Competent cells of *Saccharomyces cerevisiae* strain Y8800 (genotype *MATa trp1– 901 leu2–3,112 ura3–52 his3–200 gal4Δ gal80Δ cyh2*^*R*^*GAL1:HIS3@LYS2 GAL2:ADE2 GAL7:LacZ@met2)* were transformed with the expression vectors. At least one million colonies were harvested in YEPD medium to ensure a good coverage of all the different proteins present in the library. Finally, aliquots of the library were made and stored in glycerol stocks.

### Yeast transformation

The bait constructs cloned into the pDEST32 or pGBKT7 were transformed into the yeast strain Y8930 (genotype *MATα trp1–901 leu2–3,112 ura3–52 his3–200 gal4Δ gal80Δ cyh2*^*R*^*GAL1:HIS3@LYS2 GAL2:ADE2 GAL7:LacZ@met2)*, while the prey constructs cloned into the pDEST22 or pGADT7 were transformed into the strain Y8800 using the LiAc (Sigma-Aldrich) method (Schiestl and Gietz, 1989). Yeast transformed with the expression vectors pDEST32 and pGADT7 were plated on SC medium lacking the selective AA leucine (Leu). The same was done for the colonies transformed with the pDEST22 and pGBKT7, but in this case SC was used without the selective AA Trp. 4-day-old single colonies growing at 30°C were isolated and the insertion of the plasmids was confirmed with colony PCR using the primers listed in Supplemental Table 8. Positive transformed colonies were resuspended in YEPD containing 24% glycerol and stored at −80°C. To test for auto-activation of the bait constructs, yeast strain Y8930 carrying the expression vectors pDEST32 or pGBKT7 were grown on a SC medium lacking His and adenine (Ade) in the following combinations: -Leu -His; -Leu -His + 2 mM 3-AT; -Leu -His + 5 mM 3-AT and -Leu – Ade for pDEST32 constructs, in the case of colonies carrying the pGBKT7 the medium was lacking Trp instead of Leu. Colonies expressing the proteins PIF4 or PIF5 were able to activate the *HIS3* and *ADE2* reporter genes and for this reason they were not used in the experiments in the bait conformation.

### Y2H cDNA library screening and individual interactions

The Y2H library screening was performed using a mating-based approach described previously (Fromont-Racine et al., 2002). The yeast bait construct expressing *KDR* cloned in the pDEST32 was grown overnight in 10 ml YEPD medium under shaking conditions at room temperature. The day after a 1 ml aliquot of yeast prey cDNA library was thawed on ice, mixed with 100 ml YEPD and incubated for 1 h. The library was then pelleted at 1800 rpm for 5’ and washed two times with sterile ddH_2_O, after which it was resuspended with 10 ml YEPD. Finally, the OD_600_ of the prey library and of the yeast containing the bait construct was measured and they were mixed in equal amount of OD_600_ = 6. The yeast mix was spun down at 1800 rpm for 5’, resuspended in 300 µl of sterile ddH_2_O and plated on a 10-cm round plate containing YEPD supplemented with carbenicillin (100 µg/ml). The YEPD plate was incubated at 30°C for 4 h to allow the mating of the yeast. After the incubation, the YEPD plate was washed with 3 ml of sterile ddH_2_O, the yeast suspension was collected, centrifugated at 1800 rpm for 5’ and the pellet was resuspended in 600 µl of sterile ddH_2_O. Finally, the yeast was plated on 15-cm round plate containing SC medium – Leu -Trp -His supplemented with carbenicillin (100 µg/ml) and incubated at 30°C for 4 days. After the period of incubation, colonies growing on the selective medium were picked, resuspended in 25 µl of sterile ddH_2_O and plated on two fresh SC -Leu -Trp – His plates for 2 days at 30°C. Hereafter individual colonies of yeast from one plate were used for colony PCR using the primers listed in Supplemental Table 8. The resulting product reactions were purified using Agentcourt AMPure XP beads (Beckman Coulter) according to the manufacture’s protocol and sequenced to identify the prey proteins interacting with the bait of interest. The second -Leu -Trp -His plate was replica plated on SC -Leu -Trp -His + 2 mM 3-AT and on SC -Leu -Trp -His + 5 mM 3-AT, which were subsequently incubated for 2-3 days at 30°C, and also plated on SC -Leu -Trp -Ade for 5 days at 20°C. Yeast colonies expressing at least one of the two reporter genes were considered positive. A selection of the proteins found in the screening were full-length cloned in the prey vector, as described previously, and the yeast strain Y8800 was transformed. All the baits and the preys used for individual interactions were grown for two days at 30°C on SC -Leu or -Trp, based on the type of vectors in which they were expressed. A small dot of a single yeast colony containing the bait or the prey was resuspended in 400 µl of sterile ddH_2_O. Then 10 µl of prey yeast was mixed with other 10 µl of the bait yeast. 5 µl were spotted on YEPD plate and incubated for 24 h at 30°C to allow the mating and the growth of the yeast. Finally, a small dot of individual mated colony was resuspended in 1 ml of sterile ddH_2_O and 10 µl was spotted on the following SC selective plates: -Leu -Trp, as confirmation of the mating; -Leu -Trp -His, as confirmation of the interaction; and – Leu -Trp -His + 2 mM 3-AT; -Leu -Trp -His + 5 mM 3-AT; -Leu -Trp -Ade to determine the strength of interaction. The plate lacking Ade was incubated at 20°C for 5 days while all the others at 30°C for 2-3 days. The yeast transformed with the empty vector pDEST32 (bait) in combination with the studied preys and the empty vector pDEST22 (prey) in combination with the different baits of interest were used as negative control.

### Yeast prey plasmid TFs library of *A. thaliana*

The construction of the prey TFs library of *A. thaliana* was made as described by Pruneda-Paz et al. (2014). Briefly, the library consists of 1956 TFs cloned in full-length sequence (78.5% of all *A. thaliana* TFs) into the Gateway pDEST22 vector and expressed in the yeast strain PJ69-4A. The library is divided into 21 96-well plates and each well contains 100 µl of yeast expressing a single TF mixed to the freezing medium (SC supplemented with 22.5% glycerol) and stored at −80°C.

### Y2H TFs library screening

The bait vector pDEST32 harboring the gene *KDR* was used to transform the yeast strain PJ69-4α. The auto-activation of the bait construct was tested by taking a small dot of yeast colony, resuspended in 50 µl of sterile ddH_2_O, and spotting 5 µl on the selective plates SC -Leu -His containing 0, 5, 10, 15, 20 or 40 mM 3-AT. No auto-activation was found. From the glycerol stock, the bait yeast was grown on SC -Leu at 30°C for 2 days. At the same time, also the yeast of the TFs library was grown, the 96-well plates were thawed on ice and 5 µl were taken from each single well and spotted on a plate containing SC -Trp and left to grow at 30°C for 2 days. Then, the bait strain was resuspended in 11 ml of sterile ddH_2_O and 3 µl were spotted on YEPD plates. A small dot of each colony of the library was taken with a pipette tip, resuspended in 200 µl of sterile ddH_2_O and 3 µl were spotted on top of the bait. The plates were grown for 3 days at 30°C to allow mating and growth, after which each colony spot was resuspended in 200 µl of sterile ddH_2_O and 3 µl were plated on SC – Leu -Trp -His supplemented with 10 mM of 3-AT. The plates were incubated for 3 days at 30°C and then moved to room temperature for 3 extra days. The growth on the yeast was checked after 2, 3, 4 and 6 days to score for interactors. The identity of the positive colonies was confirmed by sequencing the result of a colony PCR used to amplify the Gateway cassette of the pDEST22 carried by the yeast. The primers listed in the Supplemental Table 8 were used to perform the colony PCR.

### Gene cloning for localization, colocalization and BiFC experiments

For *in planta* localization and colocalization experiments, cDNA deriving from *A. thaliana* was used to amplify the CDSs of *KDR, HFR1, PAR1* and *PAR2*, while AIF2, AIF4, IBH1 and IBL1 were amplified from the respective cloned in pDEST22 (previously described) using the primers listed in the Supplemental Table 9, which were designed in a way that the CDSs were in frame with a C-terminal tag and without the stop codon. The PCR products containing the *att*B sequences were cloned into the Gateway donor vector pDONR207 (gentamycin 20 µg/ml). The resulting entry vector containing *KDR* was recombined into the Gateway destination vector pEarleyGate 102 (Earley et al., 2006) (kanamycin 50 µg/ml), while *HFR1, PAR1, PAR2, AIF2, AIF4, IBH1* and *IBL1* were recombined into the Gateway destination vector pEarleyGate 101 (kanamycin 50 µg/ml).

For BiFC experiments, the following set of Gateway destination vectors were used in order to reconstitute the Venus fluorescent protein (YFP): pDEST-^GW^VYNE (N-terminal part of Venus, residues 1-173, referred to as YN in the results sections and cloned in frame to the C-terminal part of the protein of interest), pDEST-^GW^VYCE (C-terminal part of Venus, residues 156-239, referred to as YC and cloned in frame to the C-terminal part of the protein of interest), pDEST-VYNE^GW^ (N-terminal part of Venus, residues 1-173, referred to as YN and cloned in frame to the N-terminal part of the protein of interest), pDEST-VYCE^GW^ (C-terminal part of Venus, residues 156-239, referred as YC in the result section and cloned in frame to the N-terminal part of the protein of interest) (Gehl et al., 2009). Transformed competent cells were all selected based on growth on kanamycin (50 µg/ml). The proteins of this study were cloned without the stop codon when the N- or C-terminal part of the Venus protein was fused to their C-terminal part. They were cloned with the stop codon when the N- or C-terminal part of the Venus protein was fused to their N-terminal part. For each BiFC experiment, we used as positive control the combination of the two interacting proteins published with this BiFC set of vectors (Gehl et al., 2009) and as negative controls the two combinations of empty vectors with the proteins of interest.

### Transient expression in *N. benthamiana*

Competent cells of *A. tumefaciens* AGL-1 were transformed with the Gateway expression vectors described in the previous paragraph made for localization, colocalization or BiFC experiments. Transformed colonies were selected using the antibiotic resistance of the different vectors and with rifampicin (20 µg/ml) carried by AGL-1 cells. Single colonies were grown for 2 days at 28°C in 20 ml LB medium under shaking conditions. After the OD_600_ was measured, the cells were pelleted and resuspended to a final OD_600_ of 0.5 with a ½ MS medium (Duchefa Biochemie) supplemented with 10 mM MES hydrate (Sigma-Aldrich), 20 g/L sucrose (Sigma-Aldrich), 200 µM acetosyringone (Sigma-Aldrich) at pH 5.6 and incubated in darkness for at least 1 hour. The solutions were used to agroinfiltrate the abaxial side of 4-5-week-old *N. benthamiana* leaves using a 1-ml syringe without the needle. In the case of colocalization or BiFC experiments, the cells of *A. tumefaciens* carrying the two different expression vectors were mixed before performing agroinfiltration. The plants were left to grow in normal light conditions and after 2 days leaf sections were taken from the agroinfiltrated regions and visualized with a confocal microscope.

### Confocal microscopy

Microscopy was performed using a Zeiss LM 700 (Zeiss, Germany) confocal laser-scanning microscope using the 20x water immersion objective (Plan-Apochromat 20x/0.8 M27). Fresh leaf material was prepared on a glass slide with cover slip. Excitation of YFP, CFP and autofluorescence of chlorophyll was done at 488 nm, 405 nm and 488 nm. Light emission of YFP was detected at 493-550 nm, CFP at 300-483 nm and chlorophyll autofluorescence at 644-800 nm. Pinhole, gain, laser power and detector offset were always set the same within experiments. Analyses of the images were performed with ZEN lite (blue edition).

### Statistical analysis

Growth data were analyzed by 2-way ANOVA followed by post-hoc Tukey test, while qRT-PCR data were analyzed by Student’s t-test or 1-way ANOVA followed by post-hoc Tukey test. All the analyses were done using GraphPad Prism.

### Accession Numbers

Sequence data from this article can be found in the EMBL/GenBank data libraries under accession numbers: *AT1G26945* (*KDR*), *AT5G25760, AT4G26410, AT1G02340* (*HFR1*), *AT2G42870* (*PAR1*), *AT3G58850* (*PAR2*), *AT3G06590* (*AIF2*), *AT1G09250* (*AIF4*), *AT2G43060* (*IBH1*), *AT4G30410* (*IBL1*), *AT2G43010* (*PIF4*), *AT3G59060* (*PIF5*), *AT5G61270* (*PIF7*), *AT2G46970* (*PIL1*), *AT4G14130* (*XTH15*), *AT1G65310* (*XTH17*), *AT5G39860* (*PRE1*), *AT4G28720* (*YUC8*), *AT3G15540* (*IAA19*), *AT4G32280* (*IAA29*).

## ACKNOWLEDGMENTS

We would like to thank Guido van den Ackerveken for sharing the Y2H library; Mike Boxem, Manon Neilen and Marciel Pereira Mendes for help with starting Y2H library screenings; Jaime Martínez-García for sharing seeds of several mutants; Yorrit van de Kaa for help with seed propagation and screening of T_2_ and T_3_ transgenic seeds; Kaisa Kajala for help with setting up the confocal microscope for the localization experiments and Diederik Keuskamp for help with light spectral calculations.

Funding was provided by the Netherlands Organisation for Scientific Research (NWO) to Ronald Pierik (VIDI grant no. 864.12.003, Building Blocks of Life grant no. 737.016.012, VICI grant no. 865.17.002).

## AUTHOR CONTRIBUTIONS

S.B. and R.P. designed the research; S.B., C.K.P., V.H., K.v.G., and E.R. performed the experiments; S.B. and R.P. wrote the manuscript.

## Supplemental Data

**Supplemental Figure 1:**
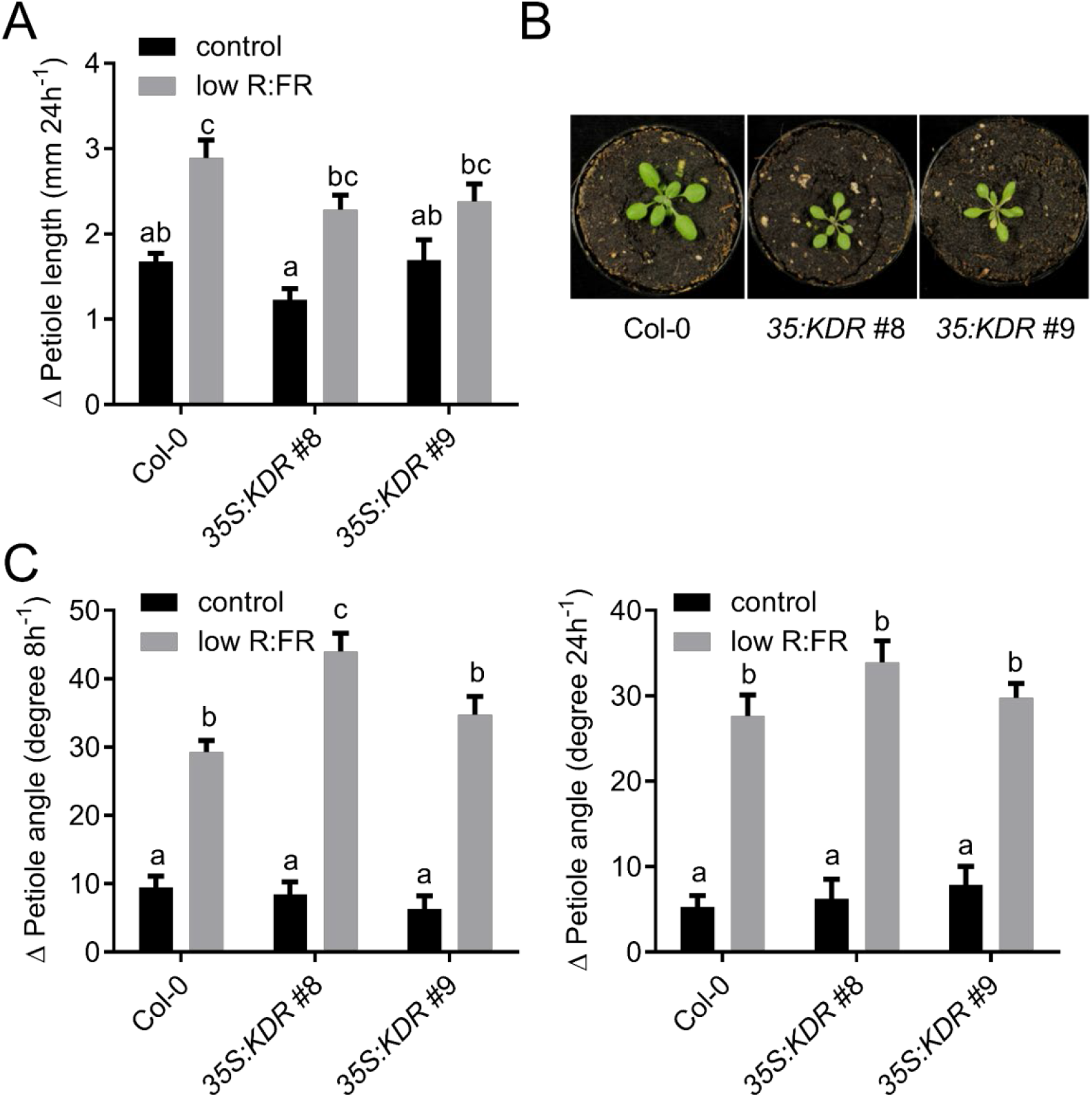
Petiole hyponasty in response to low R:FR in adult *KDR* overexpression lines. (**A**) Δ Petiole length (mm) of rosette plants of *A. thaliana* wild type (Col-0) and two independent transgenic lines overexpressing *KDR* in Col-0 background grown in control light condition (R:FR = 2) or low R:FR (R:FR = 0.2) after 24 h of light treatment. Data represent mean ± SE, n = 16. Different letters indicate statistically significant differences (2-way ANOVA with post-hoc Tukey test, p < 0.05). (**B**) Representative rosette plants of *A. thaliana* wild type (Col-0) and of two independent transgenic lines overexpressing *KDR* in Col-0 background grown for 18 days in pots containing soil in control light condition (R:FR = 2). (**C**) Differential petiole angle of rosette plants of *A. thaliana* wild type (Col-0) and two independent transgenic lines overexpressing *KDR* in Col-0 background grown in control light condition (R:FR = 2) or low R:FR (R:FR = 0.2) after 8 h (left) and 24 h (right) of light treatment. Data represent mean ± SE, n = 16. Different letters indicate statistically significant differences (2-way ANOVA with post-hoc Tukey test, p < 0.05).

**Supplemental Figure 2:**
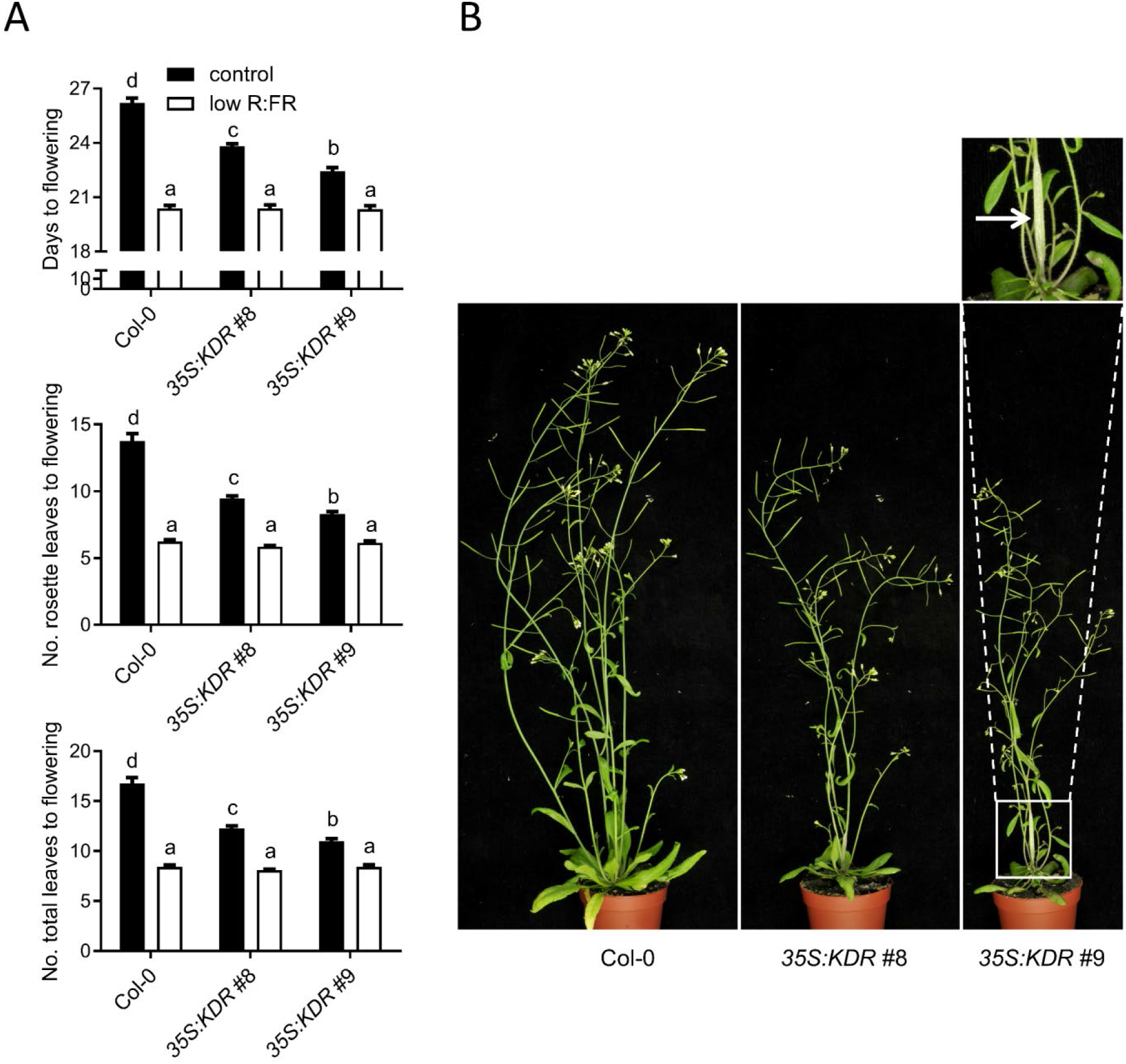
*KDR* overexpression lines show altered traits at adult stage. (**A**) Number of days to flowering (top), number of rosette leaves to flowering (middle) and total leaves number to flowering (bottom) of *A. thaliana* wild type (Col-0) and two independent transgenic lines overexpressing *KDR* in Col-0 background grown in control light condition (R:FR = 2) or low R:FR (R:FR = 0.2). Data represent mean ± SE, n = 21. Different letters indicate statistically significant differences (2-way ANOVA with post-hoc Tukey test, p < 0.05). (**B**) Representative flowering plants of *A. thaliana* wild type (Col-0) and two independent transgenic lines overexpressing *KDR* in Col-0 background grown for 35 days in pots containing soil in control light condition (R:FR = 2).

**Supplemental Figure 3:**
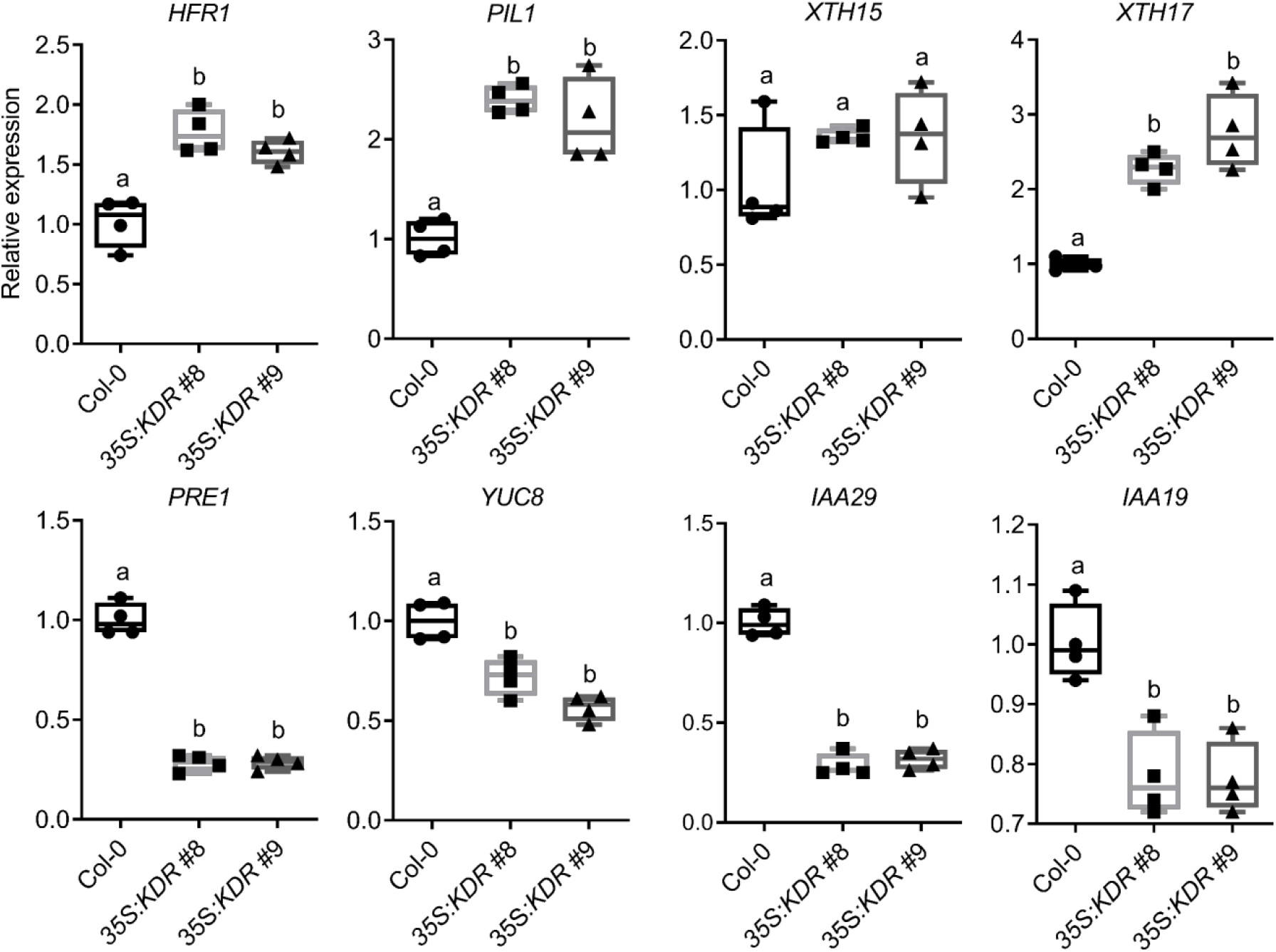
Differential regulation of PIF targets in *KDR* overexpression lines. Relative expression of known PIF targets determined by qRT-PCR in shoots of wild type Col-0 and two independent homozygous transgenic lines overexpressing *KDR*, grown in control white light condition (R:FR = 2) for 4 days. Data represent mean ± SE, n = 4. Different letters indicate statistically significant differences (1-way ANOVA with post-hoc Tukey test, p < 0.05).

**Supplemental Figure 4:**
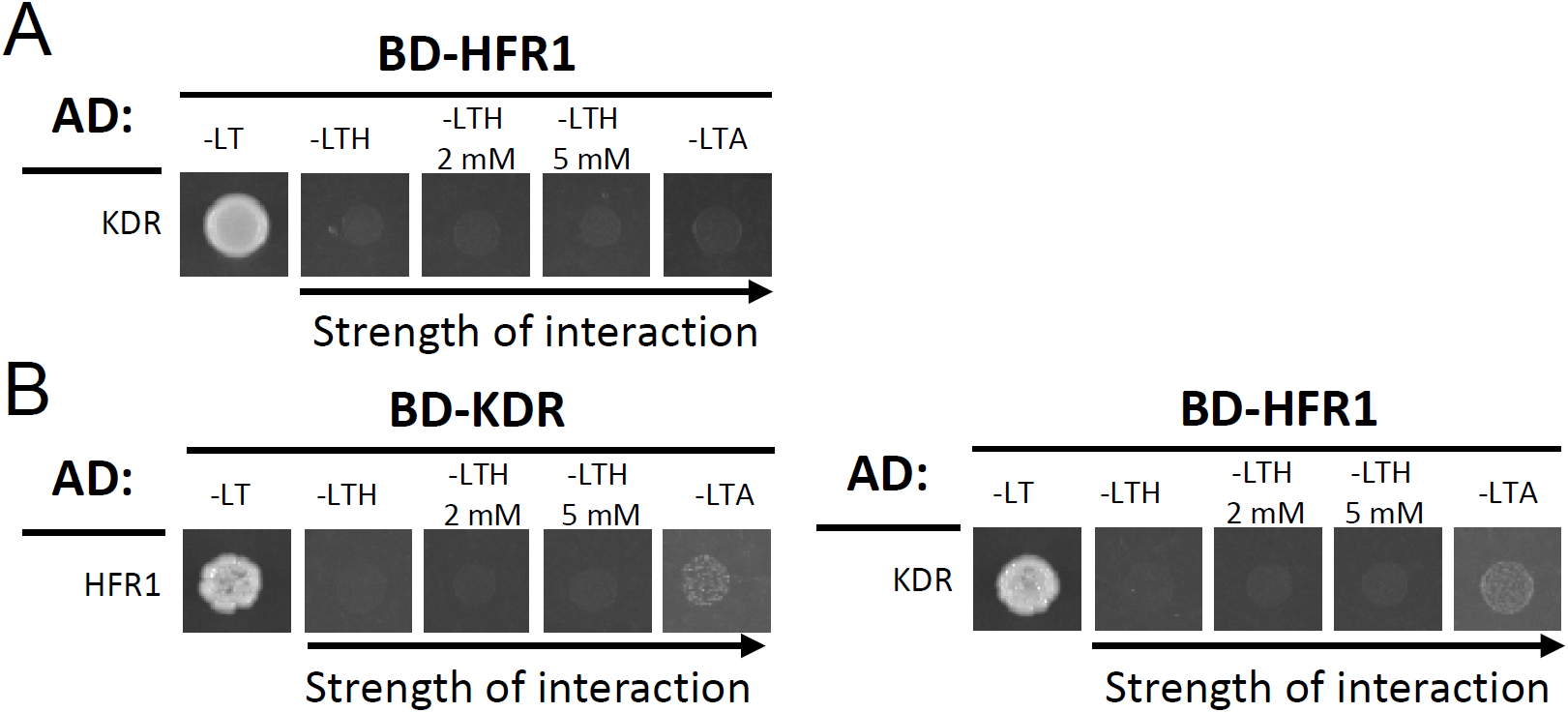
No interaction found for KDR and HFR1 in Y2H protein-protein interaction assays. (**A**) In the GAL4 Y2H assay, the GAL4 DNA-binding domain (BD) fused to HFR1 was coexpressed with the GAL4-activation domain (AD) fused to KDR. The mating of the yeast was confirmed through growth on non-selective medium (–LT). No interaction was found, as shown for lack of growth in all the different selective media (-LTH + 2 or 5 mM 3-AT and – LTA). For this experiment, the pDEST32 (bait) and pDEST22 (prey) were used. (**B**) The interaction between KDR and HFR1 was studied using KDR as bait and HFR1 as prey in the left picture. The same interaction was studied in the other conformation, as shown in the right picture. For this experiment, the vectors pGBKT7 (bait) and pGADT7 (prey) were used. No interaction was found also in this experiment. L: leucine; T: tryptophan; H: histidine; A: adenine; 3-AT: 3-amino-1, 2, 4-triazole. CFP: cyan fluorescent protein; YFP: yellow fluorescent protein.

**Supplemental Figure 5:**
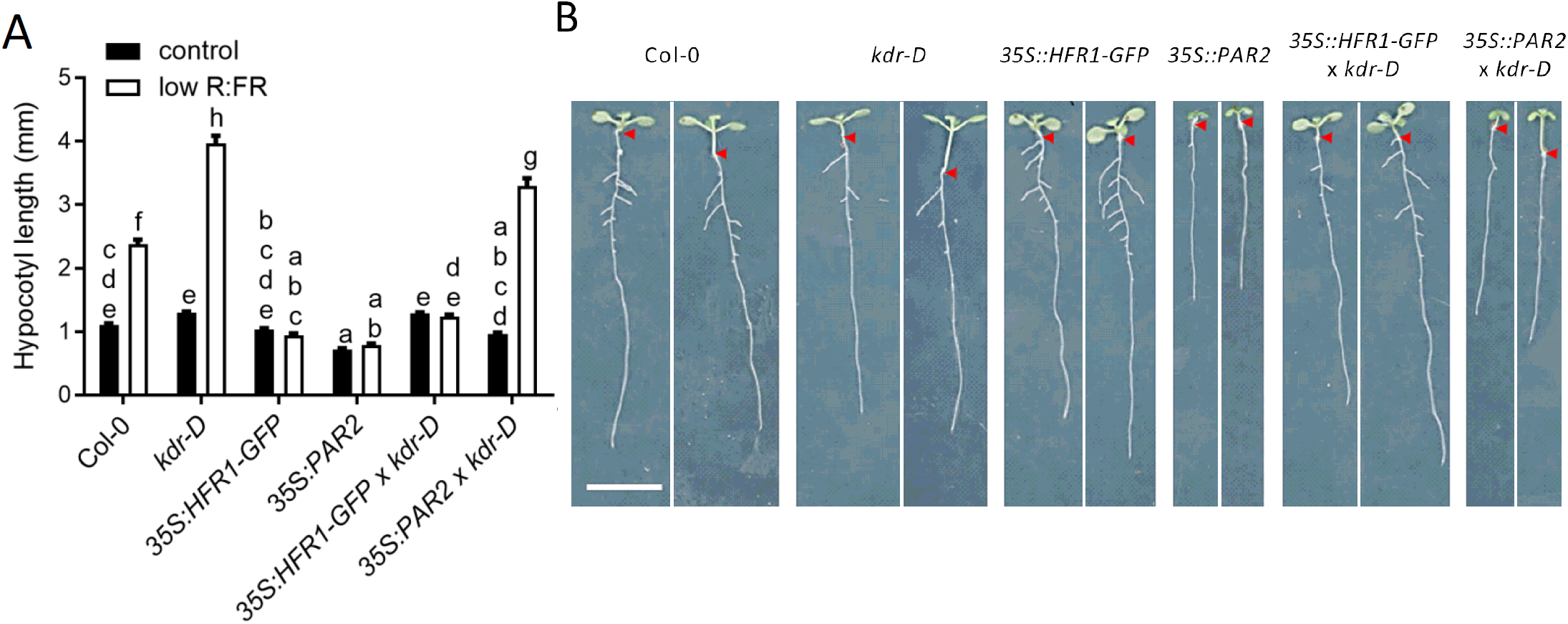
Activation-tagged *kdr-D* rescues the phenotype of *PAR2*, but not *HFR1* overexpression under low R:FR. (**A**) Hypocotyl length (mm) of seedlings of *A. thaliana* wild type (Col-0), activation-tagged line (*kdr-D*), overexpressing lines (*35S:HFR1-GFP* and *35S:PAR2*) and crossed lines of *35S:HFR1-GFP* × *kdr-D* and *35S:PAR2* × *kdr-D* grown in control light condition (R:FR = 2) or low R:FR (R:FR = 0.2) after 5 days of light treatment. Data represent mean ± SE, n = 38. Different letters indicate statistically significant differences (2-way ANOVA with post-hoc Tukey test, p < 0.05). (**B**) Representative seedlings of hypocotyl length experiment in (**A**), for each genotype the growth is shown in control light (left) and low R:FR (right). The arrows indicate the hypocotyl-root transition. The scale bar represents 1 cm.

**Supplemental Figure 6:**
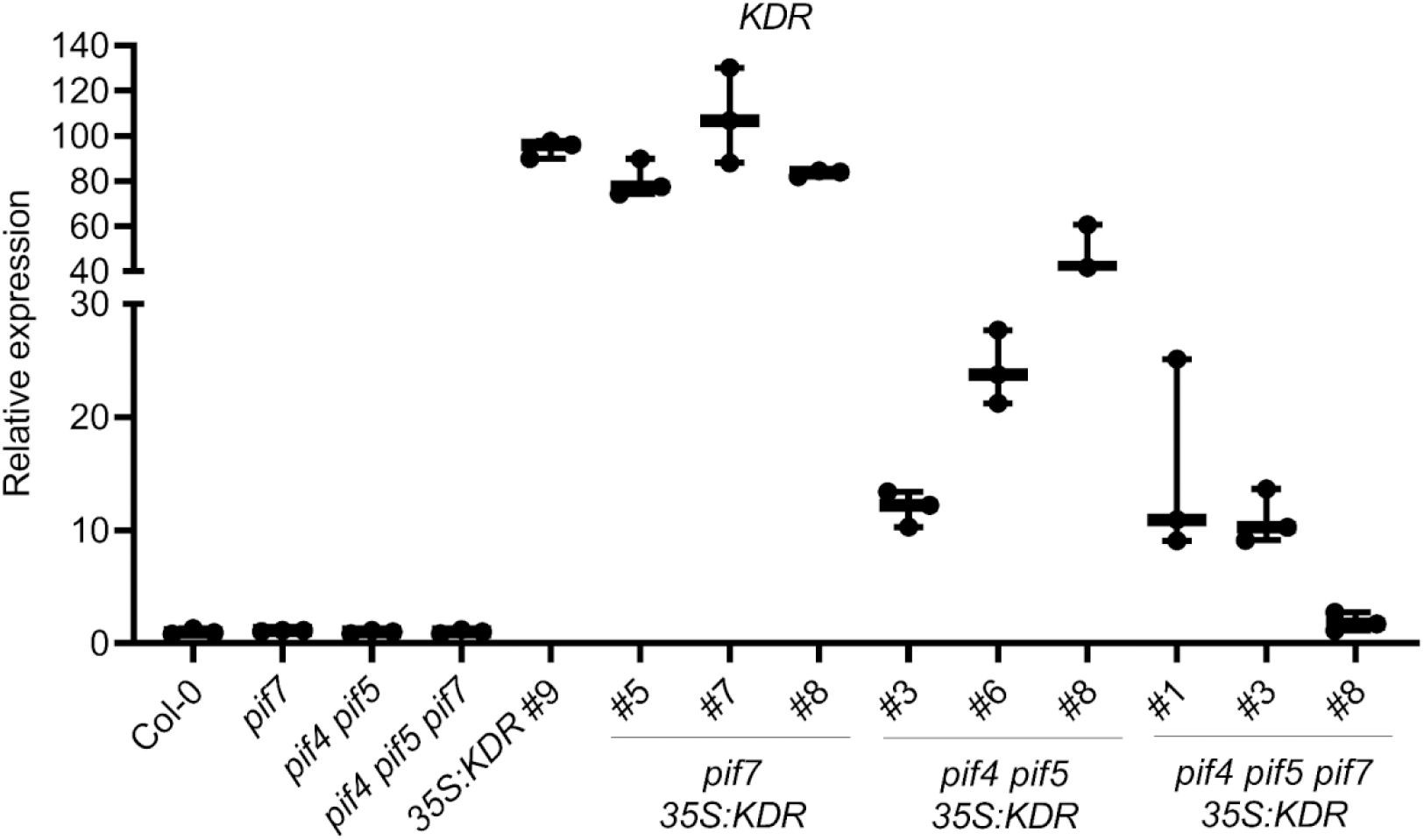
Characterization of *KDR* expression level in independent transgenic lines. Relative expression level of *KDR* determined by qRT-PCR in wild type Col-0, knockout lines (*pif7, pif4 pif5* and *pif4 pif5 pif7*), transgenic line #9 overexpressing *KDR* and independent homozygous transgenic lines overexpressing *KDR* in *pif7, pif4 pif5* and *pif4 pif5 pif7* backgrounds grown in white light. Data represent mean ± SE, n = 3.

**Supplemental Figure 7:**
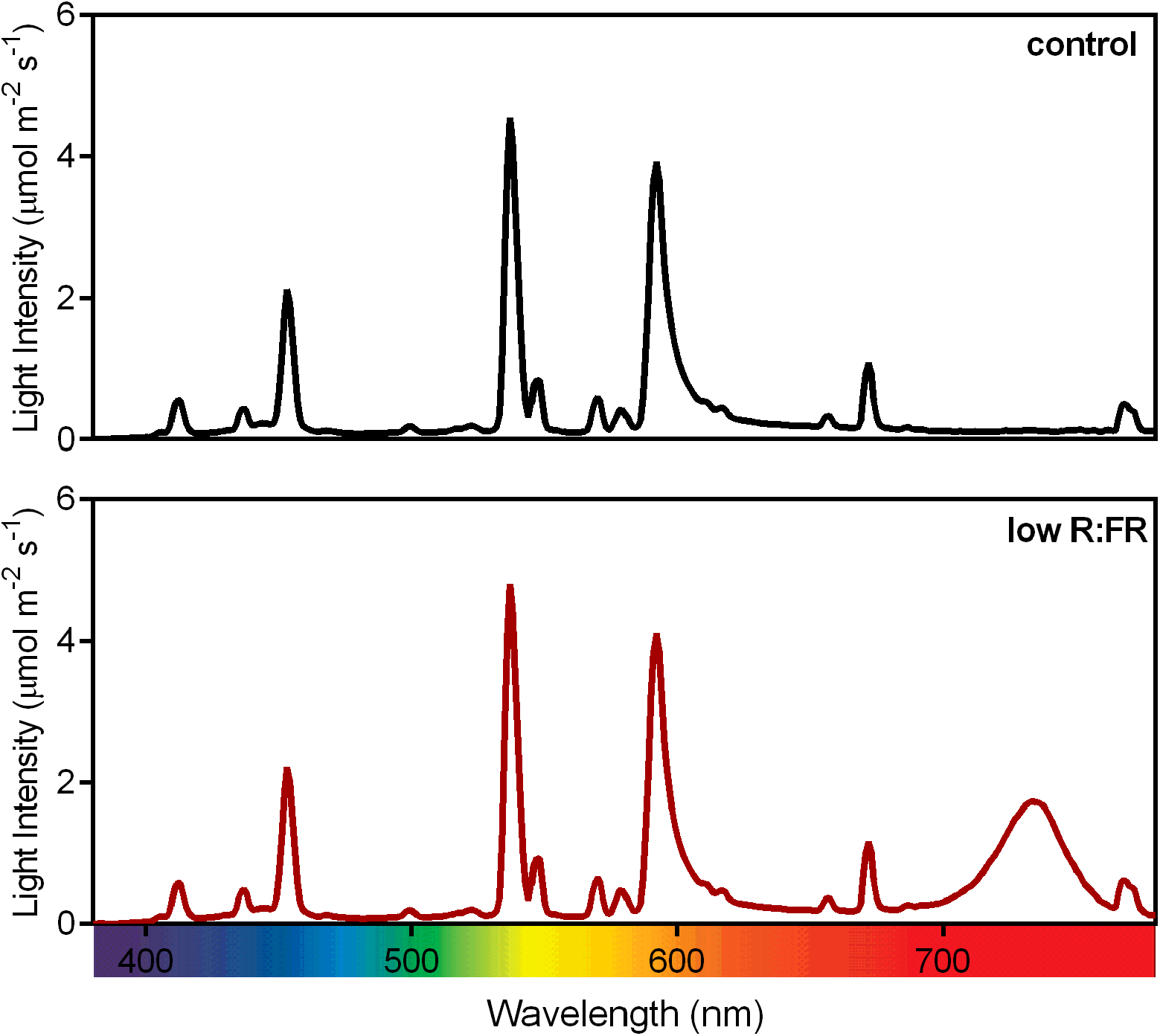
Spectral composition of the different light conditions used.

**Supplemental Figure 8:**
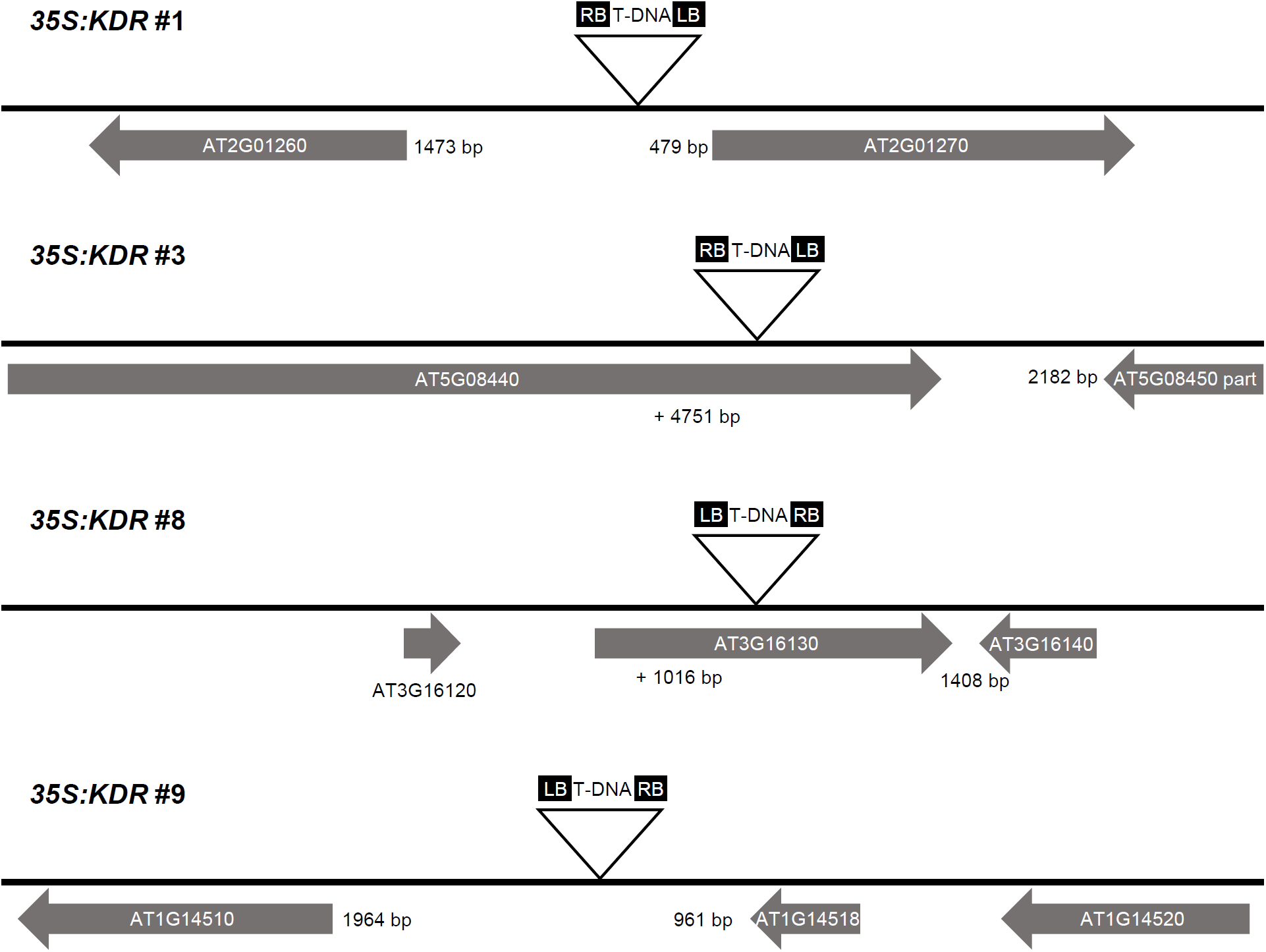
Schematic representation of T-DNA insertion sites. The insertion sites drawn with a triangle show the orientation of the T-DNA in the genome and with the numbers below is indicated the distance (bp) either from the start codon or from the stop codon relatively to the two closest genes. Genes are in scale.

**Supplemental Table 1:**
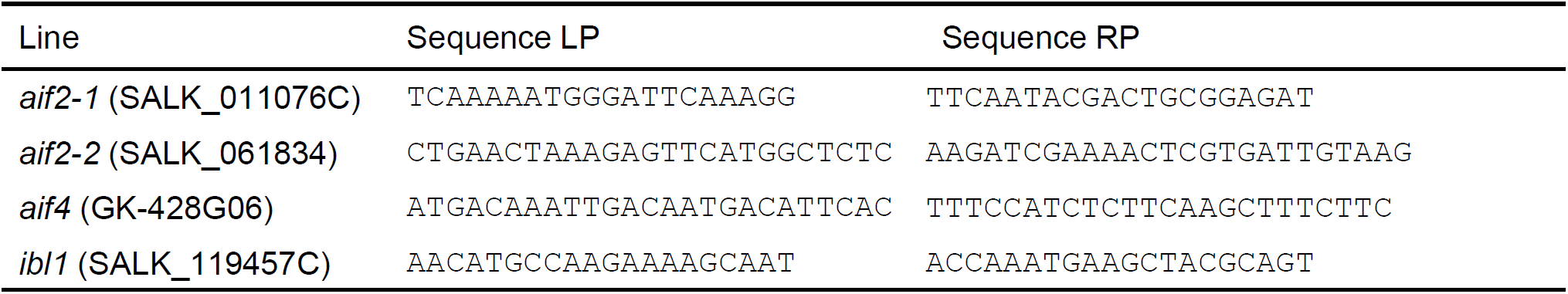
Primers used for genotyping.

**Supplemental Table 2:**
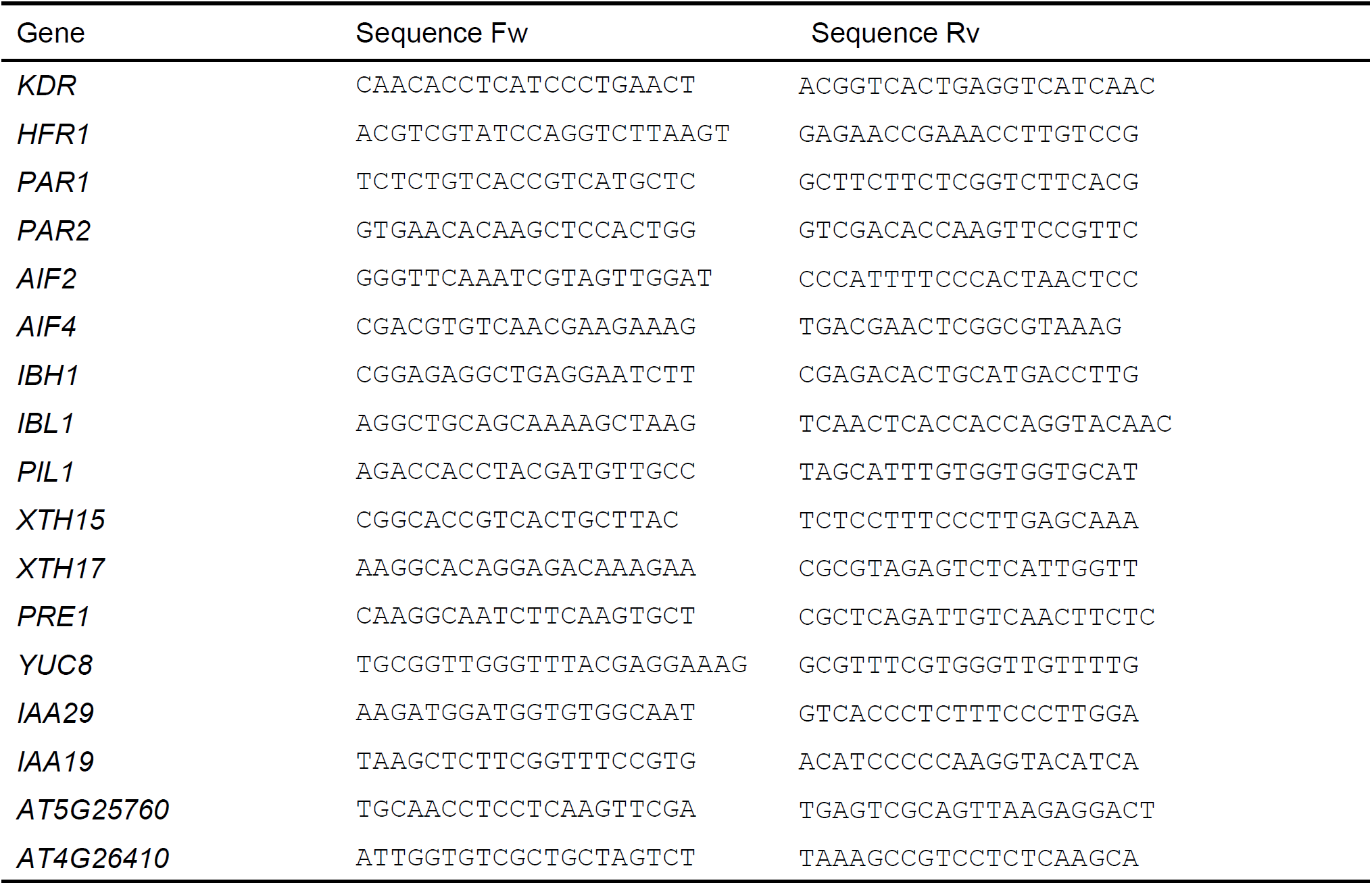
Primers for qRT-PCR.

**Supplemental Table 3:**
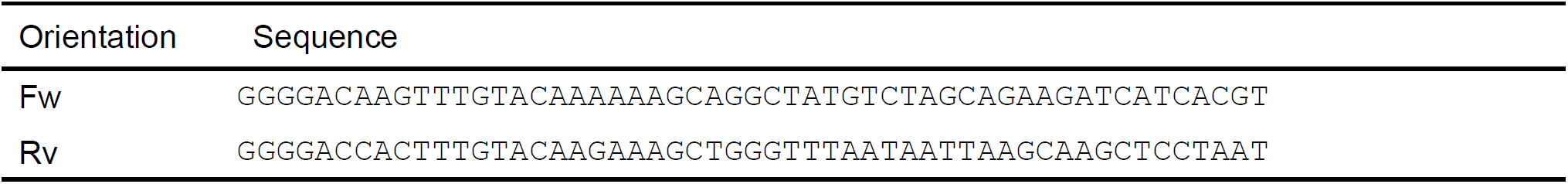
Primers with *att*B cloning sites for full-length *KDR* CDS amplification.

**Supplemental Table 4:**
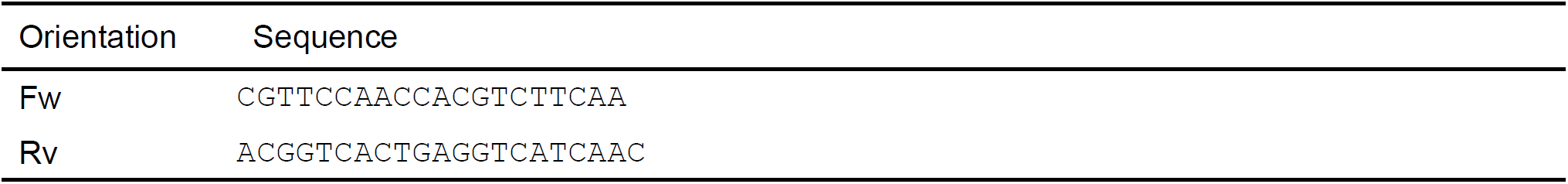
Primers used for genotyping transgenic lines overexpressing *KDR* made using the vector pFAST-G02.

**Supplemental Table 5:**
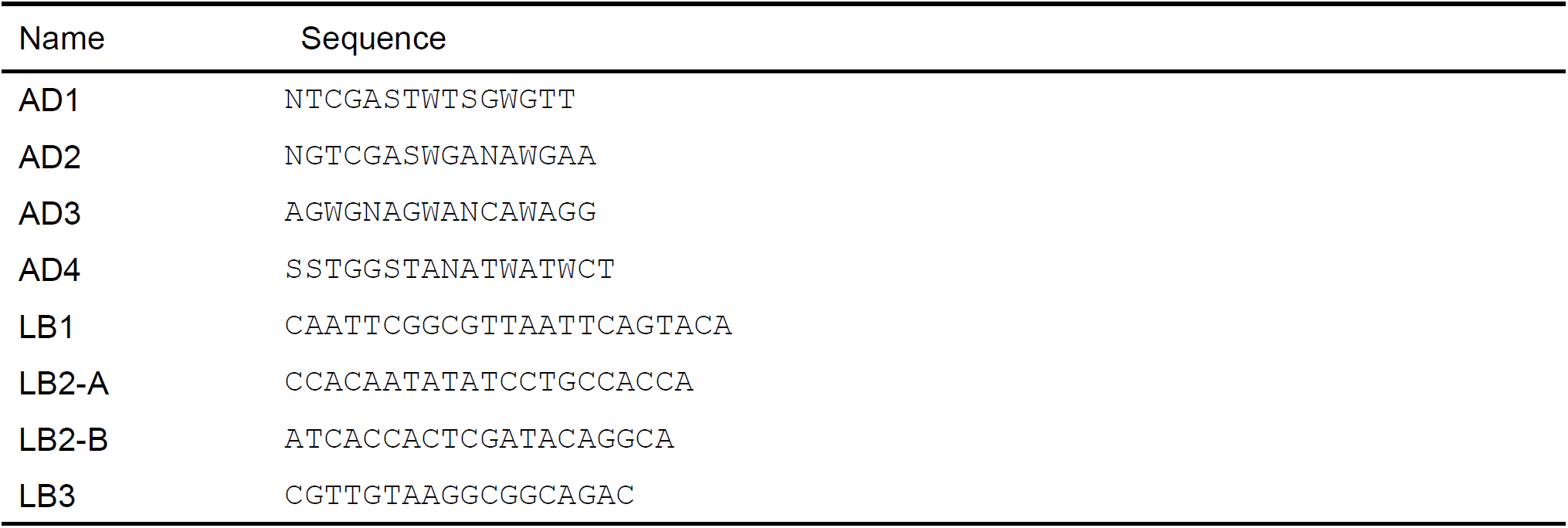
Primers used for TAIL-PCR.

**Supplemental Table 6:**
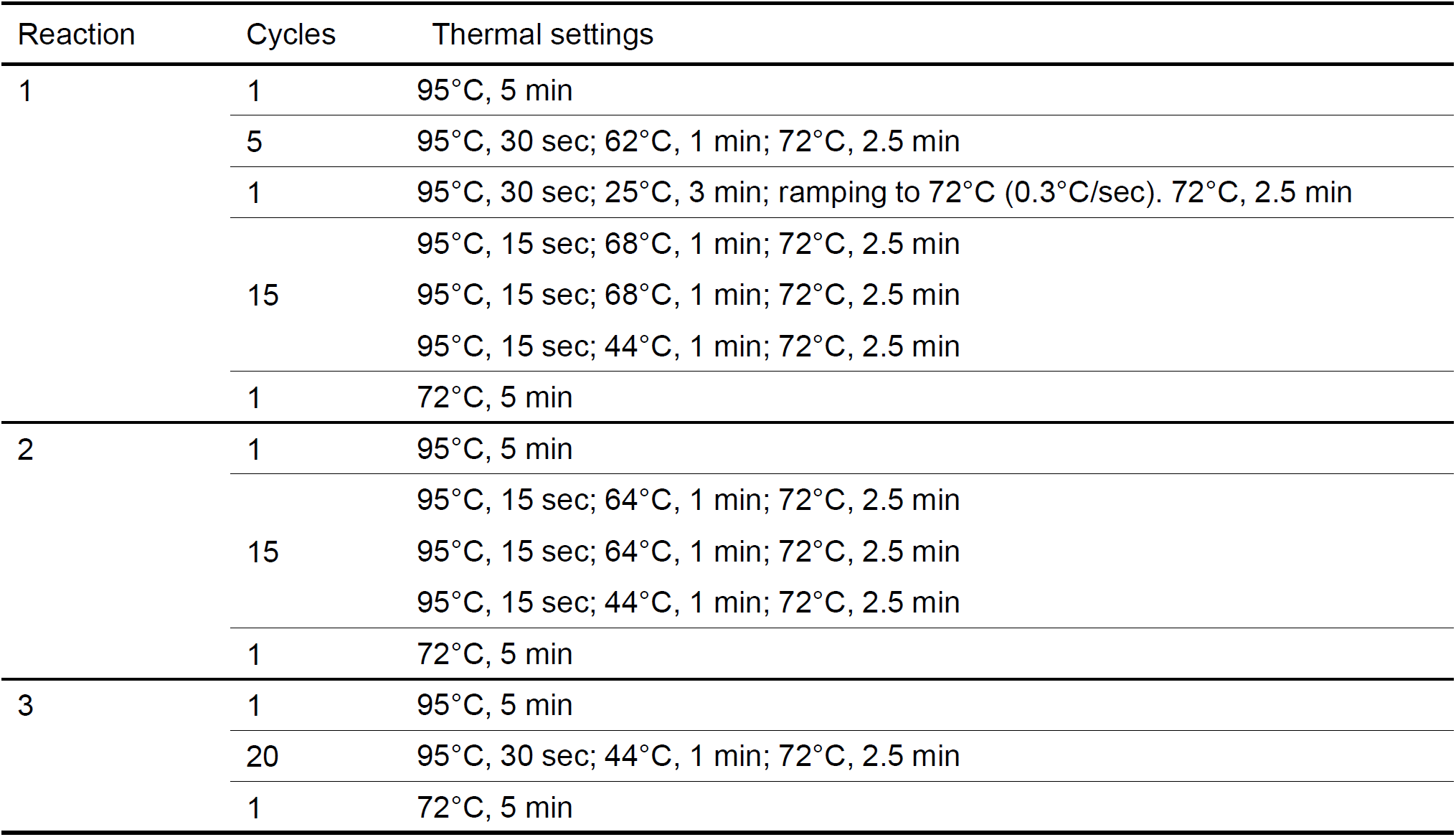
Cycle settings used for TAIL-PCR.

**Supplemental Table 7:**
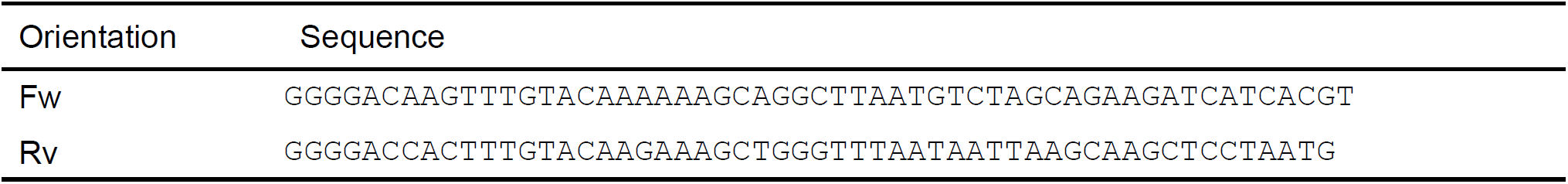
Primers with *att*B cloning sites for full-length *KDR* CDS amplification.

**Supplemental Table 8:**
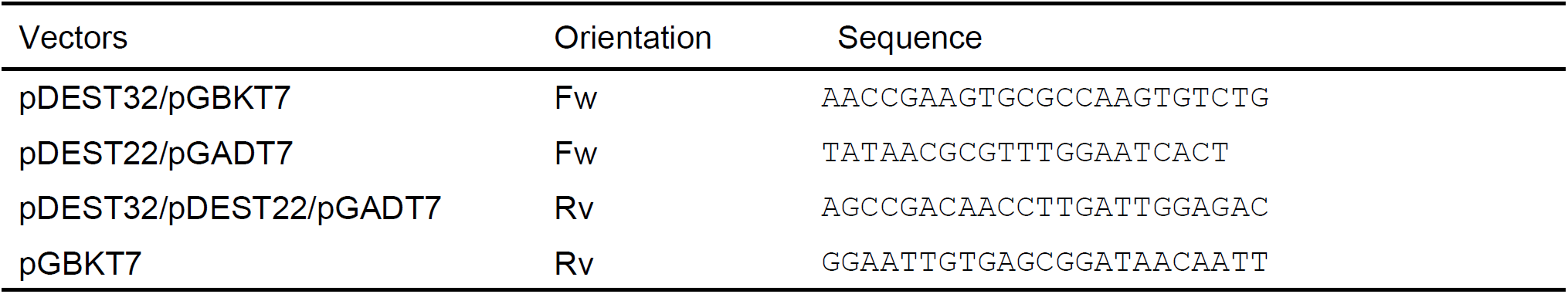
Primers used for amplification of *A. thaliana* gene fragments in Y2H vectors.

**Supplemental Table 9:**
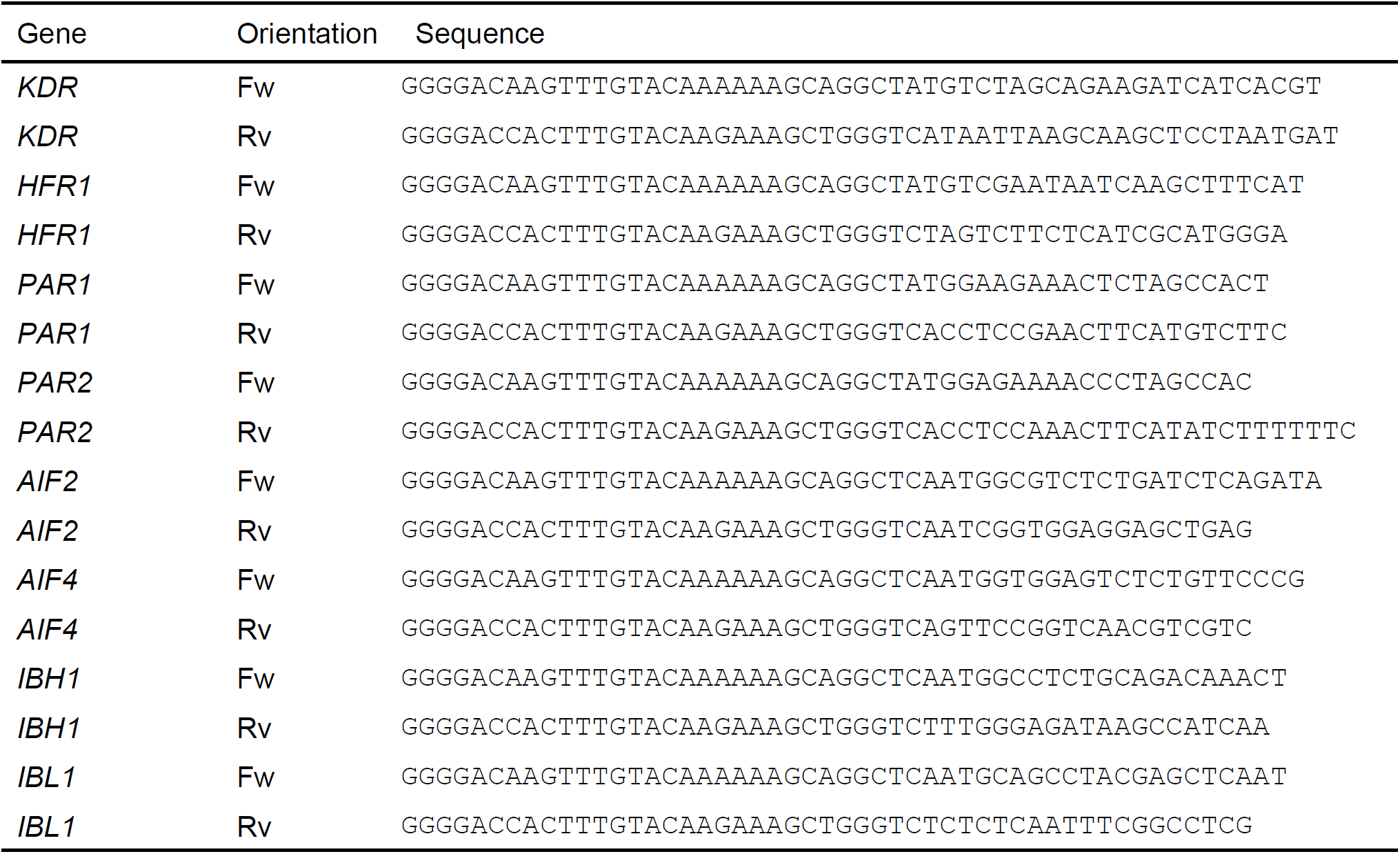
Primers with *att*B cloning sites for CDS amplification without the stop codon.

